# FtsW activity and lipid II synthesis are required for recruitment of MurJ to midcell during cell division in *Escherichia coli*

**DOI:** 10.1101/230680

**Authors:** Xiaolong Liu, Nils Y. Meiresonne, Ahmed Bouhss, Tanneke den Blaauwen

## Abstract

Peptidoglycan (PG) is the unique cell shape-determining component of the bacterial envelope, and is a key target for antibiotics. PG synthesis requires the transmembrane movement of the precursor lipid II, and MurJ has been shown to provide this activity in *E. coli.* However, how MurJ functions *in vivo* has not been reported. Here we show that MurJ localizes both in the lateral membrane and at midcell, and is recruited to midcell simultaneously with late-localizing divisome proteins and proteins MraY and MurG. MurJ septal localization is dependent on the presence of a complete and active divisome, lipid II synthesis and PBP3/FtsW activities. Inactivation of MurJ, either directly by mutation or through binding with MTSES, did not affect the midcell localization of MurJ. Our study visualizes MurJ localization *in vivo* and reveals a possible mechanism of how MurJ functions during cell division, which gives possibilities for future investigations and further antibiotics developments.

## Introduction

Rod-shaped bacteria such as the Gram-negative bacterium *Escherichia coli* grow by elongation and binary fission. One of the requirements for proliferation is the enlargement of their cell envelope, which contains an inner membrane (IM) and an outer membrane (OM). The shape of the bacterium is determined by the shape of its peptidoglycan (PG) layer, which is attached to the outer membrane in the periplasmic space located between both membranes of the envelope (Vollmer, Blanot, & De Pedro, 2008). The PG layer is a mesh-like heteropolymeric macromolecule of alternating *N*-acetylmuramyl-peptides (MurNAc-pentapeptides) and *N*-acetylglucosamine (GlcNAc) disaccharides that are connected by a β-(1,4) bond forming glycan chains that are crosslinked between *meso* diaminopimelic acid (DAP) and D-Ala at the third and fourth position of the acceptor and donor stem peptide, respectively (Bouhss, Trunkfield, Bugg, & Mengin-Lecreulx, 2008). The biosynthesis of PG is a complicated process that can be divided into 3 stages. In the cytoplasmic stage, the nucleotide precursors UDP-GlcNAc and UDP-MurNAc-pentapeptide are synthesized from fructose-6-phosphate by the GlmSMU and MurABCDEF proteins, respectively (Barreteau et al., 2008). In the IM stage, the undecaprenyl-pyrophosphate-(C_55_) linked intermediates lipid I (C_55_-MurNAc-pentapeptide) and lipid II (C_55_-MurNAc(-pentapeptide)-GlcNAc) are formed by the MraY and MurG proteins, respectively (Bouhss et al., 2008). During this stage, lipid II, the building unit of peptidoglycan, is translocated by a “flippase” across the IM (Barreteau et al., 2008; Pomorski & Menon, 2006; Ruiz, 2015, 2016; Scheffers & Tol, 2015). In the periplasmic stage, the MurNAc-pentapeptide-GlcNAc component of lipid II is inserted into the PG layer by the glycosyltransferase and transpeptidase activities of various PBPs (Macheboeuf, Contreras-Martel, Job, Dideberg, & Dessen, 2006; Sauvage, Kerff, Terrak, Ayala, & Charlier, 2008), and the lipoyl moiety C_55_-PP is cleaved off and recycled (Manat et al., 2014).

In *E. coli,* new PG-building units are inserted into the existing PG layer during length growth by a protein complex termed the elongasome, whereas during division new poles are synthesized from completely new material (Pedro, Quintela, Höltje, & Schwarz, 1997) by proteins that collectively have been termed the divisome (Den Blaauwen, De Pedro, Nguyen-Distèche, & Ayala, 2008). The elongasome is organized by the actin homolog MreB, which localizes underneath the IM in patches in a helix-like distribution. MreB is bound to the membrane by its amphipathic helix, and interacts with the bitopic membrane protein RodZ, the integral membrane proteins MreD and MreC, and the PG synthesis essential proteins RodA and PBP2 (Contreras-Martel et al., 2017; Den Blaauwen et al., 2008; Fenton, Mortaji, Lau, Rudner, & Bernhardt, 2016; Morgenstein et al., 2015; van Teeffelen et al., 2011). How the elongasome is regulated is not yet very clear whereas in contrast, cell division has been well investigated. During cell division, at least 20 proteins are recruited to midcell to form the divisome. The tubulin homologue FtsZ is attached to the IM at midcell by the anchor proteins FtsA and ZipA, and polymerizes to form the Z-ring (Blaauwen, Buddelmeijer, Aarsman, Hameete, & Nanninga, 1999; Sun & Margolin, 1998). Simultaneously, other proteins, such as the Zap proteins and FtsEX, are recruited to the ring to form the early divisome or protoring. The protoring then recruits the later divisome proteins FtsK, FtsQ, FtsL, FtsB, FtsW, FtsI (PBP3), PBP1b, FtsN (den Blaauwen, Hamoen, & Levin, 2017), and a number of hydrolases and regulatory proteins (Gray et al., 2015) to form the mature divisome (Aarsman et al., 2005). The arrival of FtsN completes the assembly of the core divisome, and signals to initiate septal PG synthesis (Gerding et al., 2009; Lutkenhaus, 2009; Weiss, 2015). The assembled divisome functions as a scaffold to initiate septal PG synthesis, OM constriction, and the cleavage of the PG layer to separate daughter cells (Gray et al., 2015). Septal PG synthesis is mainly orchestrated by PBP3 and PBP1b (Bertsche et al., 2006; Sauvage et al., 2008), but PBP1a can substitute for PBP1b (Suzuki, Nishimura, & Hirota, 1978; Yousif SY, Broome-Smith JK, 1985). OM constriction is organized by the Tol/Pal system, which forms a complex connecting IM and OM (Derouichei, Bénédetti, Lazzaroni, Lazdunski, & Lloubès, 1995; Dubuisson, Vianney, & Lazzaroni, 2002). Tightly controlled amidase activity is involved in the separation of the daughter cells (Priyadarshini, De Pedro, & Young, 2007).

Although the stages of PG synthesis are well understood, conflicting information on the flipping of lipid II across the IM has been published over the last few years. In *E. coli,* RodA and FtsW that belong to the SEDS (shape, elongation, division, sporulation) family, and MurJ that belongs to the MOP (multidrug/oligo-saccharidyl-lipid/polysaccharide) export super family (Ruiz, 2008), are reported as candidates of lipid II flippase, either functioning in elongation or division only or in both (Ikeda et al., 1989; Ruiz, 2015; Young, 2014). An alternative activity as glycosyltransferase was recently reported for RodA in *Bacillus subtilis* (Meeske et al., 2016), while the debate which protein is responsible for lipid II flipping is still going on. FtsW is an essential integral membrane cell division protein with 10 trans-membrane helixes (TMHs) and is conserved in most bacterial species that have a PG cell wall (Khattar, Begg, & Donachie, 1994). FtsW, PBP3 and PBP1b form a complex that is recruited to midcell at the later stage of divisome assembly (Goehring, Gonzalez, & Beckwith, 2006; Goehring, Gueiros-filho, & Beckwith, 2005; Leclercq et al., 2017). FtsW was shown *in vitro* to have lipid II binding and flipping activity, for which two charged residues in TMH 4, arginine 145 and lysine 153, appeared to be essential (Leclercq et al., 2017; Mohammadi et al., 2011, 2014).

MurJ is an essential IM protein that contains 14 TMHs with both C- and N-termini in the cytoplasm (Butler, Davis, Bari, Nicholson, & Ruiz, 2013; Butler, Tan, Joseph, & Ruiz, 2014). Depletion of MurJ will cause irregularly shaped cell, and finally result into cell lysis (Ruiz, 2008). Earlier functional and structural studies of MurJ reveal an outward-facing central cavity that is formed by TMHs 1, 2, 7 and 8, which contain several charged residues that are essential for MurJ function (Butler et al., 2013, 2014), while the observation of an open inward-facing conformation in the crystal structure of MurJ from *Thermosipho africanus* (MurJ_TA_) suggests alternative conformational changes of MurJ (Kuk, Mashalidis, & Lee, 2017). *In vivo* evidence favors MurJ over FtsW as the lipid II flippase, and depletion or inhibition of MurJ caused the accumulation of lipid II in cells (Qiao et al., 2017; Ruiz, 2008; Sham et al., 2014; Young, 2014), while *in vitro* flipping activity has thus far only be established for FtsW (Mohammadi et al., 2011, 2014). Interestingly, the function of MurJ in *E. coli* can be replaced by other flippases that lack sequence similarity, such as the O-antigen flippase Wzk from *E. coli,* and Amj and YtgP from other species (Elhenawy et al., 2016; Hong, Liu, & Reeves, 2018; Meeske et al., 2015; Ruiz, 2009). Two *in vitro* studies on lipid II binding of MurJ showed conflicting results, as one reported a higher lipid II binding affinity of MurJ compared with FtsW (Bolla et al., 2018), while the other one showed lipid II binding only by FtsW but not by MurJ (Leclercq et al., 2017). Although these data support MurJ to be involved in flipping lipid II, it is still not clear where and how MurJ functions in bacterial cells. In this study, by visualizing the cellular localization of MurJ with functional fluorescent protein fusions, we provide new evidence on how MurJ performs its function during PG synthesis.

## Results

### Construct a functional fluorescent protein fusion to the N-terminus of MurJ

To investigate the MurJ function and localization *in vivo,* fluorescent protein (FP) fusions to either the N-terminus or the C-terminus of MurJ with different repeats of Gly-Gly-Ser (GGS) linkers were constructed (X. Chen, Zaro, & Shen, 2013). Two low-copy-number plasmids, pTHV037 and pSAV057, were used to express the fused proteins under the control of a down-regulated *P_trcdown_* promoter (van der Ploeg, Goudelis, & den Blaauwen, 2015). Based on the physical properties of fluorescent proteins (Cranfill et al., 2016; Meiresonne, van der Ploeg, Hink, & den Blaauwen, 2017), green fluorescent protein mNeonGreen (mNG) and red fluorescent protein mCherry (mCh) were chosen for MurJ fusions. The functionality of the MurJ fusions was tested by a complementation assay (Material and Methods).

As shown in Table 1, the N-terminal fusions of MurJ were fully functional, while C-terminal fusions were less functional or not functional at all. Among these, the N-terminal mNG fusion with a (GGS)_2_ linker showed the most stable and strongest fluorescence signals. For mCh, the fusion without linker was the best. Thus they were chosen for further studies. The crystal structure of MurJ from *T. africanus* reveals that TMHs 13 and 14 form a hydrophobic groove, which penetrates into the central transport cavity and is hypothesized to be important for lipid II binding (Kuk et al., 2017). The loss function of MurJ C-terminal fusions might due to the defects in folding and groove formation.

**Table 1.**
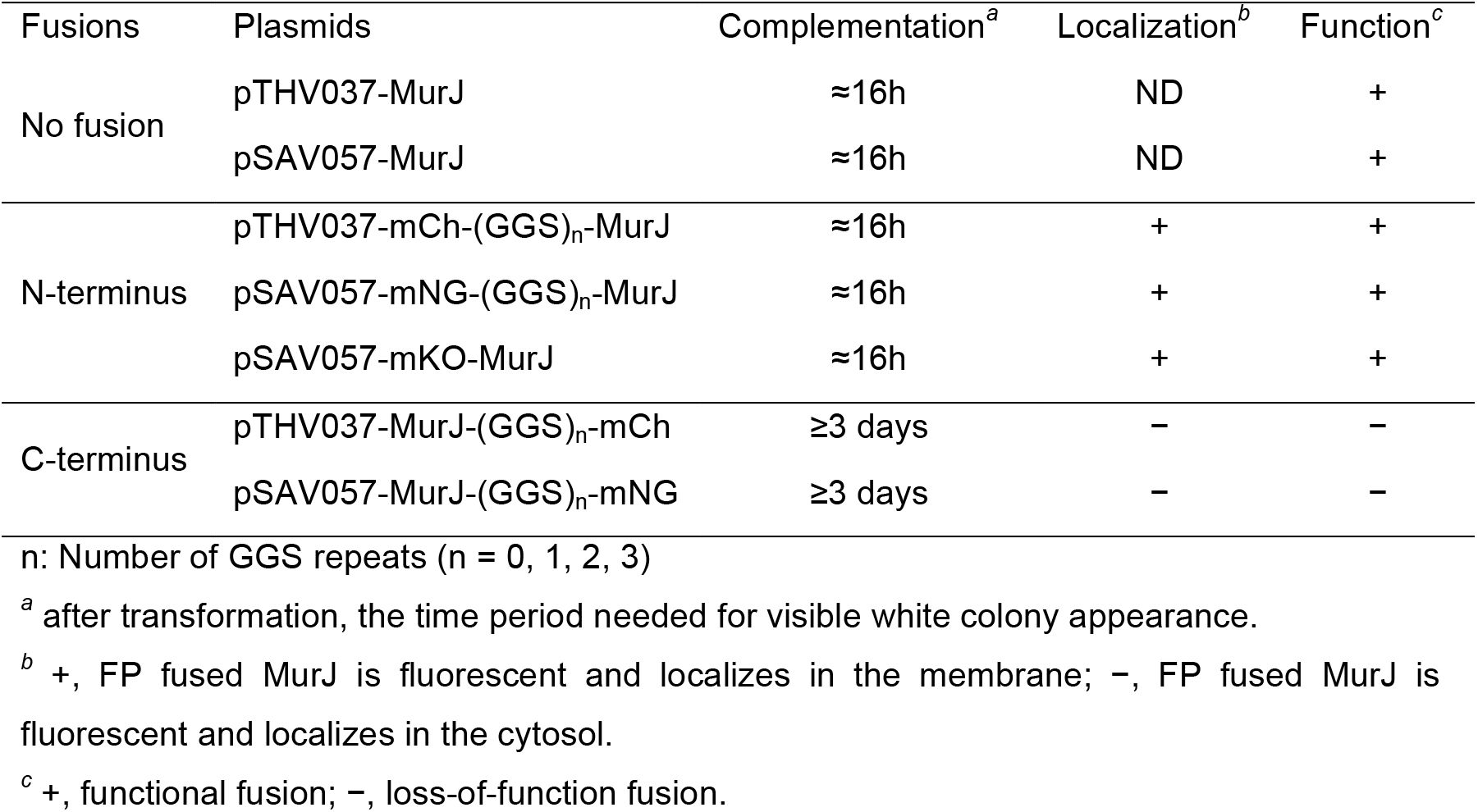
Functionality and localization of MurJ fluorescence proteins fusions with different linkers at N- or C-terminus

### MurJ localizes in the lateral wall and also at midcell

MurJ is reported to be present at 674 and 78 molecules per cell when grown in rich and minimal medium, respectively (Li, Burkhardt, Gross, & Weissman, 2014). In order to characterize MurJ at its endogenous expression level, the MurJ fusion was introduced into the *E. coli* chromosome at its native locus. The existence of the native promoter *pmurJ* was confirmed by inserting a divergently transcribed chloramphenicol cassette between the stop codon of gene *yceM* upstream of *murJ* and the putative *murJ* promoter (Fig. S1A). However, attempts to introduce FP-fused MurJ under the native *murJ* promoter failed, apart from a mCh fusion, which yielded a strain with defects in cell morphology and growth (Fig. S1B). Since the plasmid expressed fusions were fully functional, a likely explanation for the morphological defect could be that the translation efficiency of the fused proteins was less than that of MurJ only. Consequently, the expression of the FP fused MurJ under the native *murJ* promoter was not enough to support growth. Therefore, we replaced the chromosomal *murJ* gene with *fp-murJ* constructs transcribed under the control of *p_trcdown_* promoter to generate strains XL08 and XL09 (Material and Methods). No defect on growth or cell morphology was observed even when grown without IPTG induction (Fig. S1 C and D).

Strain XL08 was grown to steady state in minimal glucose medium (Gb1) at 28°C to be able to correlate its length to its cell division cycle age (Blaauwen et al., 1999; Vischer et al., 2015), and the localization of MurJ in living *E. coli* cells was analyzed using ImageJ and the Coli-Inspector of ObjectJ (Vischer et al., 2015). To avoid overexpression, no IPTG induction was applied. The mass double time (MD) of this strain is 83 minutes, which is close to 80 minutes of wild-type *E. coli* (Vischer et al., 2015). MurJ localized in the cylindrical part of the cells as well as specifically at midcell in dividing cells (Fig. 1). After normalizing and plotting the fluorescence profiles into 10 age groups, a clear midcell localization was observed starting from 50% of the cell cycle. The septal proportion of MurJ, which is indicated as the “Ring fraction”, was analyzed as function of the division cell cycle, and a maximal proportion of 8.58% was observed at 82.5% of cell cycle (Fig. S2). To show that this accumulation was not an effect of double membrane formation during cell invagination, the localization of mNG-(GGS)_2_-GlpT was introduced as control. GlpT (glycerol-3-phosphate transporter) is the major transporter for the uptake of *sn-* glycerol 3-phosphate in *E. coli.* It was reported to have 12 TMHs and to function as monomer (Huang, 2003), which diffuses homogenously in the IM (Oswald, Varadarajan, Lill, Peterman, & Bollen, 2016). It was therefore expected to show homogenous membrane localization. Like for MurJ, the localization of mNG-(GGS)_2_-GlpT was imaged and analyzed. As expected, GlpT localized homogenously in the membrane without midcell accumulation. After plotting into 10 age groups, GlpT showed a clear difference with MurJ, and no enhanced fluorescence was observed at midcell during constriction (Fig. 1A and B). The differences between the localization patterns of the two proteins are also obvious in images of the cells (Fig. 1C). The specific localization pattern of MurJ suggests that MurJ is likely involved in both length growth and cell division.

**Fig. 1.**
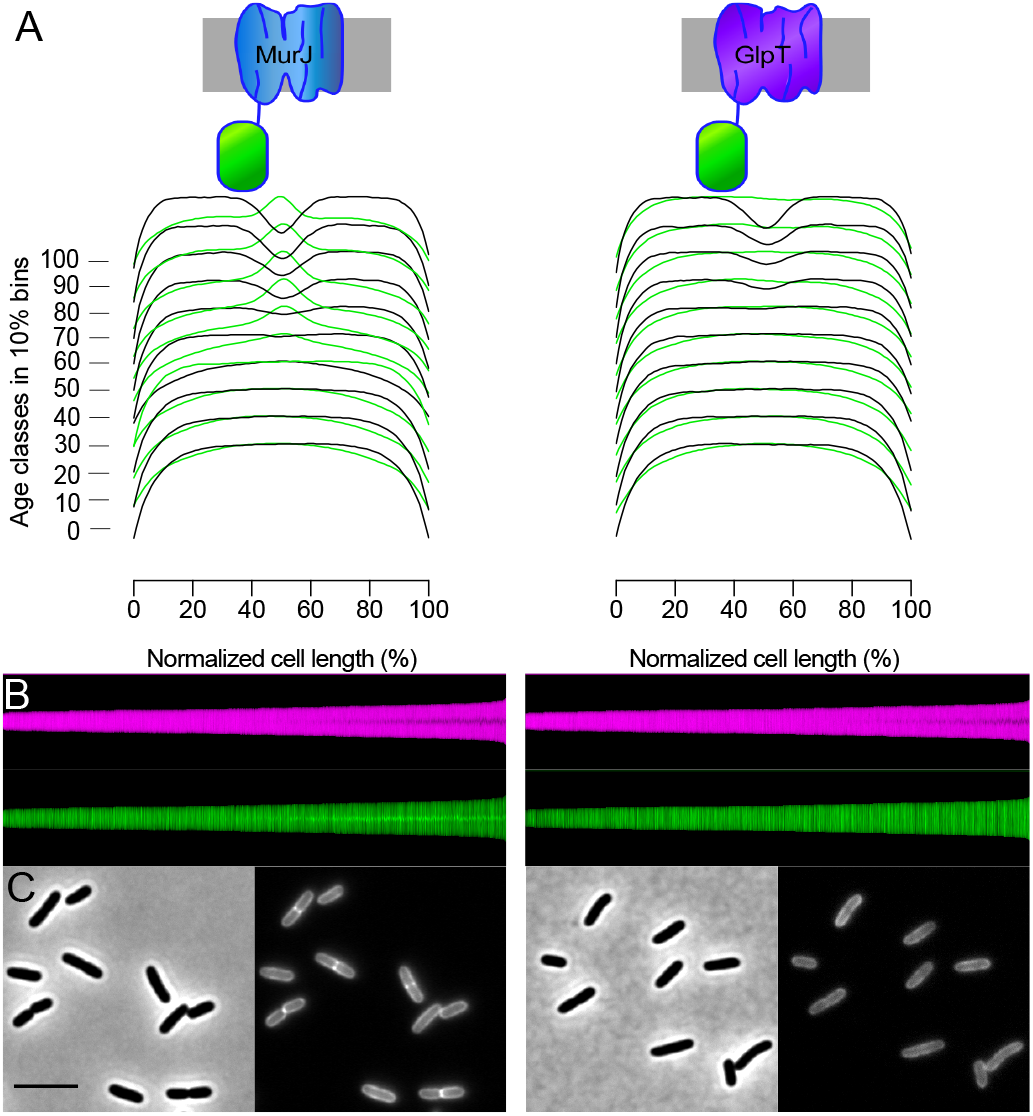
MurJ localizes both in the lateral wall and at midcell. Strain XL08 containing the chromosomal mNG-MurJ fusion shows a cell-cycle-dependent localization in contrast to the homogenously localizing integral membrane protein mNG-GlpT. Cells were grown in Gb1 minimal medium to steady state at 28°C. Left: MurJ profiles; Right: membrane control protein GlpT profiles. (A) Collective profiles of localization of MurJ and GlpT. For each protein, the diameter (black lines) and fluorescence (green lines) profiles along normalized cell length are shown in 10 % age class bins. (B) Maps of diameter profiles (magenta) and fluorescence profiles (green). Cells are plotted by length, ascending from left to right. (C) Phase contrast image (left) and the corresponding fluorescence image (right) for each protein. Scale bar equals 5 μm. More than 1200 cells were included for both analyses.

### MurJ is recruited to midcell simultaneously with the PG synthesis proteins

The observation of MurJ midcell localization prompted us to the next question: at which point of the cell division cycle MurJ is recruited. To reveal this, localization of MurJ, the early and late localizing proteins FtsZ and FtsN, respectively, and the proteins MraY and MurG, which synthesize Lipid I and lipid II, respectively (Aarsman et al., 2005; Bisson Filho et al., 2017; Bouhss, Mengin-Lecreulx, Le Beller, & Van Heijenoort, 1999; Mengin-Lecreulx, Texier, Rousseau, & Van Heijenoort, 1991; Mohammadi et al., 2007; Weiss, 2015) was determined in steady-state grown strain XL08. Cells from the same culture and optical density were either imaged live for MurJ localization or were fixed with FAGA (2.8% formaldehyde and 0.04% glutaraldehyde) and subsequently immunolabeled with antibodies specific for the above mentioned proteins (Buddelmeijer, Aarsman, & Blaauwen, 2013). The early divisome protein FtsZ was recruited at midcell at about 25% of cell cycle, and the core divisome assembly was completed by the midcell arrival of FtsN at 40%-50% of the cell division cycle (Fig.2). The measured extra fluorescence at mid cell (FCplus) of these proteins plotted as function of the cell division cycle age (Fig. S3) is in agreement with a previous study (Van der Ploeg et al., 2013). Interestingly, MraY, MurG and MurJ accumulated at midcell simultaneously at approximately 50% of cell cycle at the later stage of divisome assembly. These results indicate that MurJ is very likely functioning together with the PG synthesis complex, which would be in agreement with its function as a lipid II flippase.

### MurJ requires a mature divisome for midcell localization

The timing of MurJ midcell localization raised the question whether its localization is dependent on particular divisome proteins. To answer this, the mNG-(GGS)_2_-MurJ chromosomal fusion was introduced into a series of temperature-sensitive cell division mutant strains. FtsA is an essential early divisome protein that anchors the Z-ring to the IM (Pichoff & Lutkenhaus, 2005; Rico, García-Ovalle, Mingorance, & Vicente, 2004). The absence of FtsA causes filamentous growth of *E. coli*, as the downstream divisome proteins are not recruited to midcell (Pichoff & Lutkenhaus, 2002). In the FtsA(*Ts*) background, MurJ showed normal midcell localization when grown at permissive temperature. However, when grown at 42 °C, MurJ midcell localization was lost, and MurJ only localized in the cylindrical membrane of the filamentous cells (Fig. 3B and Fig. S4). This indicates that MurJ midcell localization is dependent on the presence of the early divisome. Like in the FtsA(*Ts*) cells, depletion of the later divisome proteins, FtsQ, FtsW, FtsI or FtsN (Ishino et al., 1989; Taschner, Huls, Pas, & Woldringh, 1988), abolished MurJ midcell localization in the filamentous cells (Fig. 3 C-F, and Fig. S4). To point out, some localization bands of MurJ were observed in FtsQ(*Ts*) filaments at 42 °C, which was likely caused by the relocalization of FtsQ proteins during the imaging at room temperature. As control, MurJ still localized at midcell in a wild-type background at 42 °C (Fig. 3A). These results suggest that MurJ requires a mature divisome for midcell localization. To determine to what extent MurJ is dependent on cell division, we also examined MurJ localization in strains that were deleted for PBP1a, PBP1b, Pal, TolA, or the amidases AmiABC. The observation of a clear MurJ midcell localization in the absence of these proteins, even when cells grow as chains, indicates that MurJ midcell localization is independent of these proteins (Fig. 3 G-K). All these results are consistent with the function of MurJ as the lipid II flippase during PG synthesis.

**Fig. 3.**
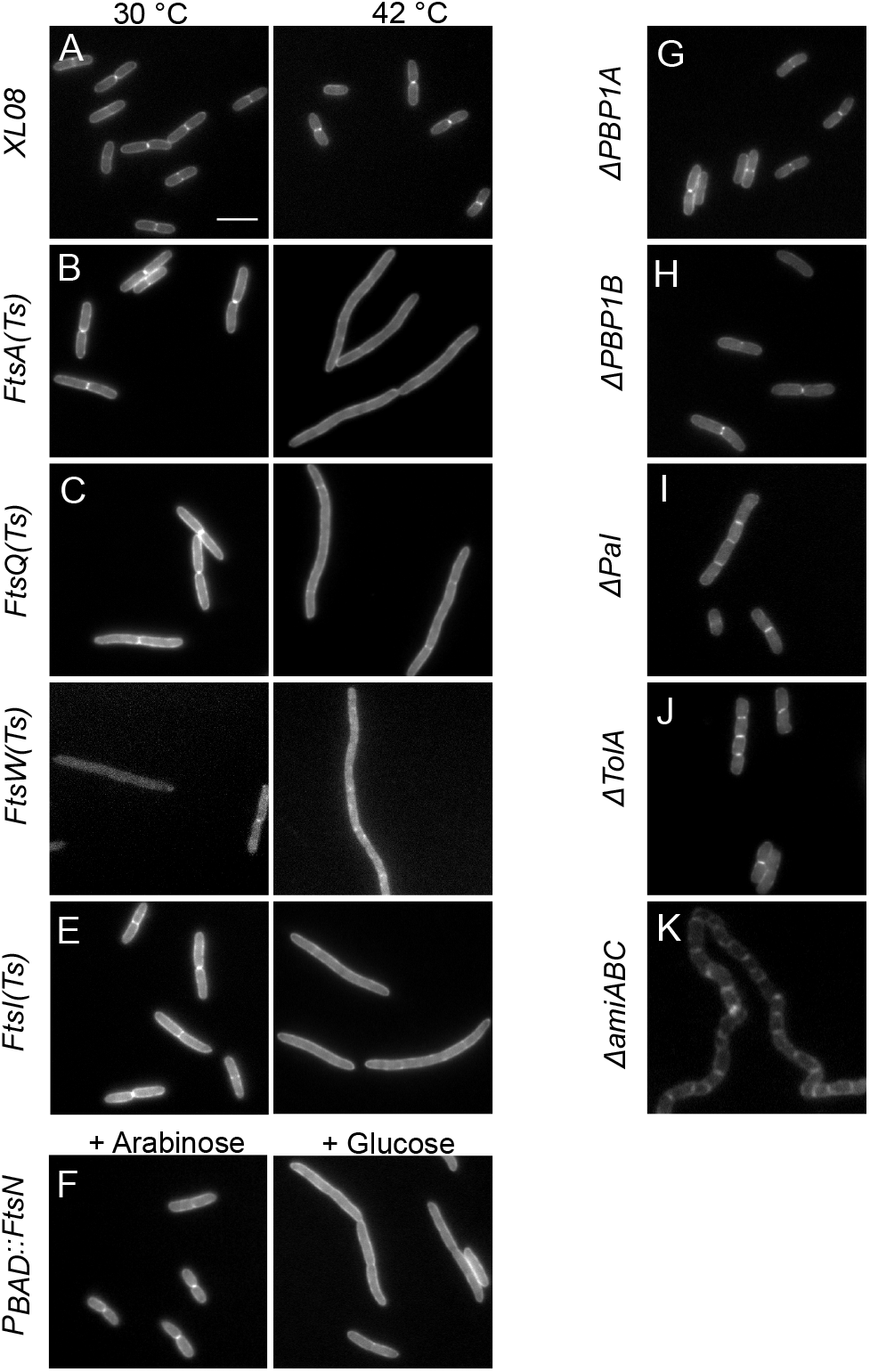
MurJ requires the core divisome for its midcell localization. Chromosomal mNG-(GGS)_2_-MurJ fusion was introduced into strains that harbor temperature sensitive, depletable, or deleted division proteins (except that the mNG-(GGS)_2_-MurJ plasmid was expressed in *ΔtolA and Δpal strains, and the* mCherry-MurJ plasmid was expressed in *ΔamiABC* strain). (A-E) MurJ localization in XL08 (WT), *ftsA 1882* (Ts), *ftsQ1* (Ts), *ftsW* (Ts) and *ftsI2185* (Ts) backgrounds. Left: Localization at permissive temperature of 28 °C. Right: Localization after growth at the non-permissive temperature of 42 °C for 2 mass doublings (MD)s. (F) MurJ localization in FtsN depletion background at 30 °C. Left: localization in the presence of 0.2% w/v arabinose. Right: Localization of MurJ in the absence of arabinose for 2 MDs. (G-H) mCh-MurJ localization in *ΔmrcA* (PBP1A), *ΔmrcB* (PBP1B), *Δpal, ΔtolA* and *ΔamiABC* background at 30 °C, respectively. Scale bar equals 5 μm.

### MurJ midcell localization requires PBP3 activity

After observation of MurJ midcell localization, the next obvious question would be whether this localization is dependent on septal PG synthesis. Septal PG synthesis requires the activity of PBP3 (FtsI) and PBP1b, although the latter can be replaced by PBP1a (Yousif SY, Broome-Smith JK, 1985). The specific inhibition of PBP3 activity by aztreonam blocks septal PG synthesis without initially inhibiting the localization of the divisome, resulting in filamentous cells with divisomes localized at regular cell division distances (Den Blaauwen, Aarsman, Vischer, & Nanninga, 2003; Mohammadi et al., 2007). After growing the chromosomal mNG-MurJ containing strain XL08 at 28 °C to steady state in Gb1 medium, MurJ localization was determined in the presence of 1 mg.L^−1^ aztreonam. A block of MurJ midcell localization was already observed after 30 min, without much change in cell morphology (Fig. S5). After 1 MD and 2 MDs, MurJ midcell localization was completely abolished in the typical filamentous cells (Fig. S5 and Fig. 4). Subsequently, cells were fixed and immunolabeled with antibodies against FtsZ and FtsN to show the localization of divisomes, and against MraY and MurG to show the localization of the lipid II-synthesizing complex. As expected, we found that FtsZ, FtsN, MurG and MraY localized at potential division sites in the filaments (Fig. 4). Likewise, MurJ localization was determined in the presence of elongasome inhibitors A22 and mecillinam for 2 MDs, which inactivate MreB and PBP2, respectively (Den Blaauwen et al., 2003; Shih, Kawagishi, & Rothfield, 2005; Vollmer et al., 2005; White & Gober, 2012). No influence on MurJ midcell localization was observed although cells started to grow spherically (Fig. S6). Together, these results suggest that MurJ localization is specifically dependent on the activity of PBP3 and septal PG synthesis.

**Fig. 4.**
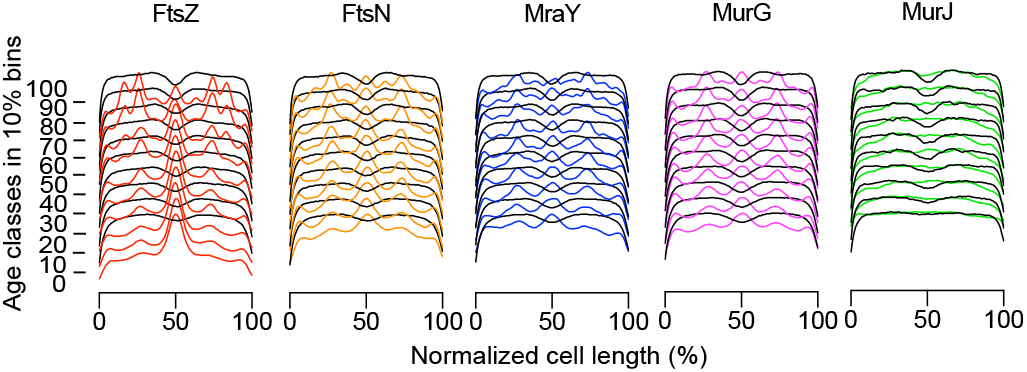
Localization of cell division proteins in the presence of the PBP3 inhibitor aztreonam for 2 MDs. Cell division is inhibited but the localization and assemblage of the divisome continues, whereas the localization of MurJ is lost. In Gb1 at 28 °C, steady state growing XL08 was treated with 1 mg.L^−1^ aztreonam, MurJ localization was determined in living cells, and FtsZ, FtsN, MraY and MurG were immunolabeled with antibodies after fixation. Diameter (black lines) and fluorescence (colored lines) profiles were plotted into 10 % age class bins along the normalized cell length. More than 1200 cells were included.

### MurJ midcell localization requires lipid II synthesis

Next, we asked whether MurJ localization would be lipid II-dependent. The biosynthesis of lipid II requires the activity of two essential enzymes in the cytoplasmic stage, the pyridoxal 5’-phosphate (PLP) dependent alanine racemase, Alr, which converts L-alanine into D-alanine (Barreteau et al., 2008; Wood & Gunsalusa, 1959), and the ATP-dependent D-ala:D-ala ligase, Ddl, which catalyzes the formation of D-alanyl-D-alanine from two molecules of D-alanine (Barreteau et al., 2008; Neuhaus, 1962). D-cycloserine (DCS) is a structural analogue of D-alanine that inactivates Alr and Ddl, and causes rapid cell lysis (Azam & Jayaram, 2015; Neuhaus & Lynch, 1972; Prosser & de Carvalho, 2013; Wang, Walsh, & Walsh, 1978). A concentration of 1 μg.ml^−1^ DCS is sufficient to deplete the UDP-MurNac-pentapeptide pool in 30 minutes when *E. coli* is grown in minimal medium (de Roubin, Mengin-Lecreulx, & van Heijenoort, 1992; Mengin-lecreulx et al., 1983). Therefore, steady-state grown XL08 cells were treated with different concentrations of DCS, and MurJ localization was determined at 30 min after adding DCS. At concentrations of 0.5 μg.mL^−1^ DCS or higher, growth inhibition and rapid cell lysis was observed after 30 min, while lower concentrations of DCS caused lysis after 1 MD or even longer incubation periods (Fig. 5A). Interestingly, DCS also showed a concentration-dependent effect on MurJ localization. Increasing DCS concentrations eventually blocked the MurJ midcell localization, and when the DCS concentration was higher than 0.5 μg.mL^−1^, MurJ was mostly absent from midcell (Fig. 5B and Fig. S7A). In addition, localization of MraY and MurG was somewhat reduces at midcell in the presence of DCS (Fig. S7, B and C). On the contrary, the core divisome monitored by the presence of FtsN remained intact (Fig. S7D). Together, these results indicate that MurJ midcell localization is dependent on the synthesis of lipid II.

**Fig. 5.**
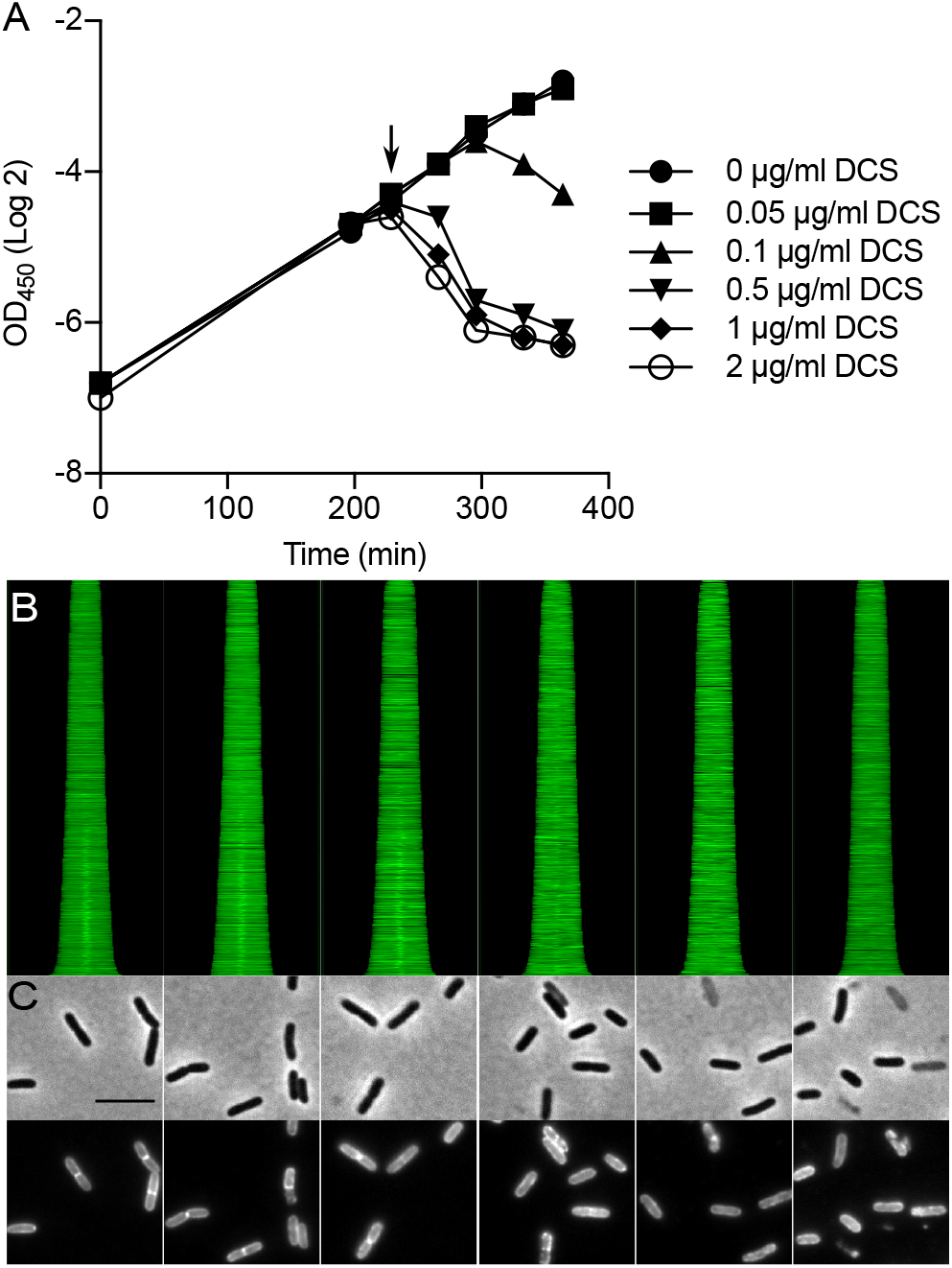
Inhibition of lipid II biogenesis blocks MurJ midcell localization. In Gb1 at 28 °C, steady state growing XL08 was treated with increasing concentrations of D-cycloserine (DCS), and mNG-MurJ localization was determined at 30 minutes after addition of DCS. (A) Growth curves were plotted as time against the natural logarithm of the optical density of the cells. The black arrow indicates the addition of DCS. (B) Fluorescence maps profiles of MurJ localization. Cells are plotted according to increasing cell length from top to bottom. From left to right, the maps correspond to 0, 0.05, 0.1, 0.5, 1 and 2 μg.L^−1^ of DCS concentrations. More than 1200 cells were included. (C) Phase contrast images (upper) and correspond fluorescence images (down) of cells treated with DCS. Scale bar equals 5 μm.

### MurJ midcell localization requires FtsW activity

Our results so far showed that MurJ midcell localization is dependent on the assembly of the divisome, the activity of PBP3 and also the synthesis of lipid II. Previous investigation of divisome assembly and regulation suggests that FtsQLB normally keeps FtsW/PBP3 inactive, and once the divisome is completely assembled, the FtsW/PBP3 complex is somehow activated (Liu, Persons, Lee, & de Boer, 2015; Weiss, 2015). Very recently, *in vitro* data showed that FtsW, PBP3 and PBP1b form a ternary complex, and the lipid II binding activity of FtsW inhibits the polymerization of lipid II by PBP1b in the absence of PBP3, while the presence of PBP3 stimulates the release of lipid II from FtsW, and activates its polymerization by PBP1b (Leclercq et al., 2017). These observations raised the possibility that FtsW is needed to ensure the lipid II accessibility to MurJ, which implied that MurJ would not be able to localize at midcell in the presence of inactive FtsW. To validate this hypothesis, plasmids that expressed two non-functional FtsW mutants, FtsW R145A and FtsW K153N were introduced into strain XL08 to investigate the localization of MurJ. These two mutants were shown to localize at midcell but failed to complement the FtsW(Ts) strain at non-permissive temperature, and their expression in LMC500 wild-type cells showed a dominant-negative filamentous morphology (Mohammadi et al., 2014). In addition, *in vitro* data showed that these mutants lost the lipid II “flipping” activity of FtsW (Mohammadi et al., 2014). In our study, to eliminate the role of the wild-type FtsW, 20 μM IPTG was used to overexpress these two mutants, and thus diluted the functional wild type FtsW that expressed from the XL08 genome. When cells were grown to steady state in minimal medium, cell length increased due to the defect on cell division, especially in cells expressing mutant K153N (Fig. 6B). MurJ midcell localization was lost in the presence of non-functional FtsW mutants for 2 MDs, despite the overexpression of MurJ as it is also expressed under the IPTG inducible *P_trcdown_* promoter. In contrast, the divisome, which was monitored by immunolabeling of FtsN, was still localized at midcell (Fig. 6). Midcell localization of MurJ was not affected when wild type FtsW was overexpressed (Fig. S8), indicating that MurJ needs a functional FtsW for its localization at mid cell. Moreover, when grown in LB medium, strains expressing these mutants showed strong defects on morphology and MurJ localization. At early exponential phase (OD_600_ = 0.2), filamentous morphology was observed for all cultures, even for cells grown with glucose that suppressed the expression of the FtsW mutants (Fig. S9A). Interestingly, in cultures at late log phase expressing the FtsW K153N mutant or in cultures at stationary phase expressing the FtsW R145A mutant, some cells started to restore the MurJ localization, especially in the cells that were induced with IPTG (Fig. S9A). Since these mutants are toxic, we wondered whether the XL08 strain lost these FtsW expressing plasmids at high frequency, restoring MurJ localization. A spot assay was carried out to test our hypothesis: cells from each culture were diluted to the same OD600 value and spotted on LB and LB-ampicillin agar plates (these two plasmids expressing the FtsW mutants are ampicillin resistant). For both strains, no obvious growth difference was observed on both plates when grown with glucose, indicating that most cells still contained the mutant plasmids. However, for cells that had been induced with IPTG, a large population of cells had lost the FtsW mutant plasmids, as much less growth was observed on the ampicillin plate compared with LB plate (Fig. S9B). These results strongly suggest that MurJ midcell localization requires FtsW activity, rather than its physical presence.

**Fig. 6.**
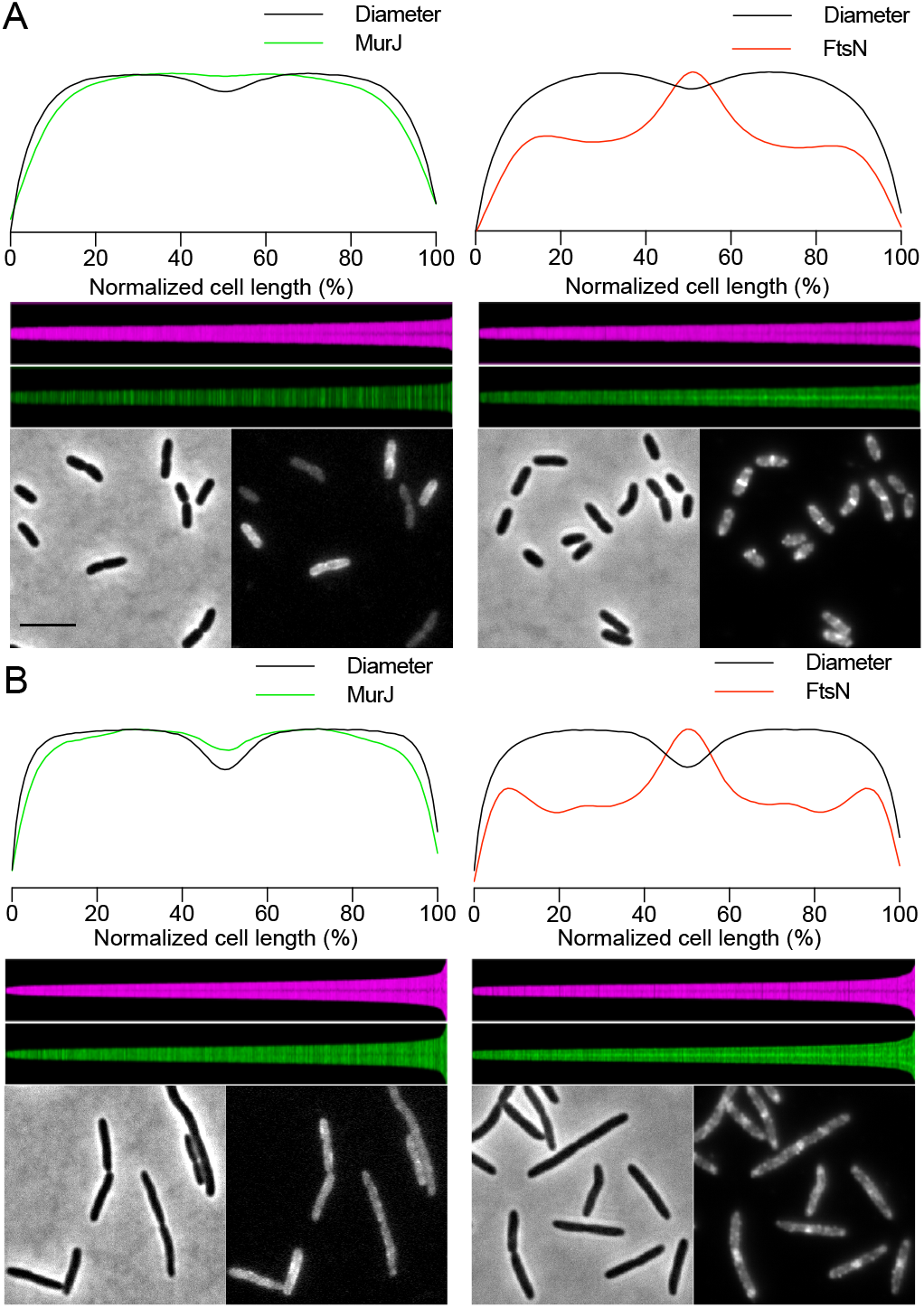
Expression of non-functional FtsW mutants abolishes MurJ midcell localization. XL08 expressing mutant FtsW R145A or FtsW K153N was grown in Gb1 minimal medium to steady state at 28 °C. IPTG (20 μM) was used to induce expression of the FtsW mutants from plasmid. MurJ localization was determined in living cells and the presence of the divisome was confirmed by immunolabeling of FtsN, after fixation. (A) Localization of MurJ and FtsN after expression of FtsW R145A. (B) Localization of MurJ and FtsN after expression of FtsW K153N. For each part: top, average localization profiles plotted along normalized cell length (n ≥ 1200 cells). Black lines indicate cell diameter. Green lines and red lines indicate MurJ and FtsN localization, respectively. Middle, map profiles of cell diameter and MurJ or FtsN localization sorted by ascending cell length. Bottom, phase contrast and fluorescence microscopy images. Scale bar equals 5 μm.

### MurJ activity is not required for its localization

Earlier studies revealed several conserved charged residues, R18, R24, R52 and R270 in the central cavity of MurJ that are essential for its function (41–43). To verify whether MurJ needed to be functional to allow midcell localization, an mNG fused MurJ R18A mutant was expressed from plasmid in LMC500. The cells were grown in Gb1 and expression was induced with 20 μM IPTG. As shown in Fig. 7A, the non-functional mutant R18A localized both in the lateral wall and at midcell as was observed for the wild type-MurJ fusion. Although MurJ is likely functional as a monomer based on its structure, the wild-type copy of MurJ from LMC500 genome might still contribute to the localization of this non-functional mutant. To eliminate this potential contribution, localization of the R18A mutant was also determined in strain XL30, a MurJ-depletion strain in which native MurJ production is under an arabinose-inducible promoter (Ruiz, 2008). In the presence of arabinose, no growth defect was observed, and the R18A mutant showed the same localization as in the LMC500 strain (Fig. 7B, left). When grown in the presence of glucose, native MurJ was eventually depleted as morphology defects were observed after 3 hours, however, the mutant R18A was still localized in lateral wall and at midcell (Fig. 7B, right). The localization of the other loss-of-function-mutants showed the same results as R18A (see below). These results suggest that the MurJ function is not required for its localization.

**Fig. 7.**
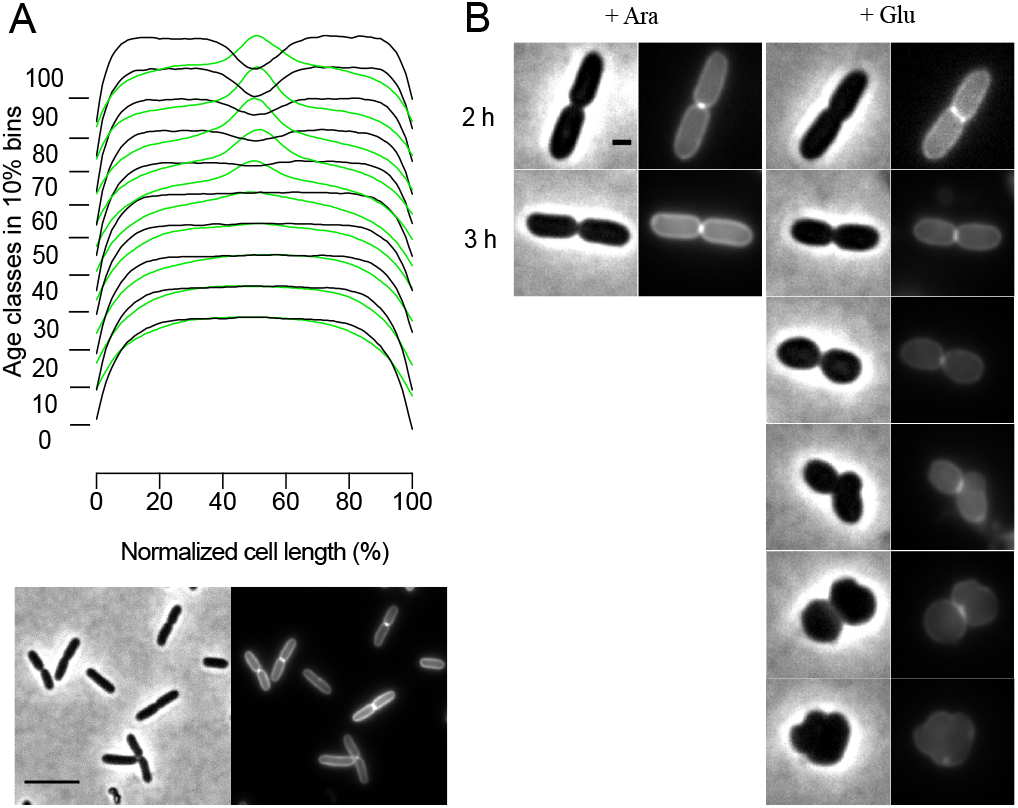
MurJ non-functional mutant R18A does not influence its localization. (A) The plasmid expressing the mNG-MurJ R18A fusion was introduced into LMC500. This strain was grown in Gb1 minimal medium to steady state at 28 °C, and expression was induced with 20 μM IPTG for 2 MDs. The diameter (black lines) and fluorescence (green lines) profiles along normalized cell length are shown in 10 % age class bins. More than 1200 cells were included. Scale bar equals 5 μm. (B) The plasmid expressing the mNG-MurJ R18A fusion was introduced into the MurJ depletion strain XL20. Mutant localization was determined in the presence of arabinose (wild type MurJ expression) or glucose (wild type MurJ depletion). Scale bar equals 1 μm.

### MTSES does not influence the localization of MurJ single-cysteine variants

The function of a number of MurJ single-cysteine mutants, A29C, N49C, S263C and E273C, was reported to be sensitive (only partially in the case of E273C) to MTSES (Butler et al., 2013). MTSES covalently modifies reduced cysteine residues that are exposed to the periplasm. In the absence of MTSES, these mutant proteins are functional, but not when MTSES is bound to their cysteine residues (Chamakura et al., 2017; Qiao et al., 2017; Sham et al., 2014). To investigate whether MTSES affects the functionality of these mutants through the disruption of their recruitment, the localization of their mNG fusions was determined in the presence of MTSES. Having optimized the induction conditions, MurJ cysteine mutants were induced with 40 μM IPTG, and their localization was determined at 10 min and 40 min (20 min for S263C variant due to the higher sensitivity) after addition of 0.5 mM MTSES (Material and Methods and Fig. S10). We found that MTSES influences only the functionality but not the localization of these mutants, as MurJ midcell localization was still observed, in spite of cell lysis after MTSES addition (Fig. 8 A-D, and Fig. S11). As a control, neither the functionality nor localization of cysteine free variant (MurJ Cys^−^) was influenced by MTSES (Fig. 8 E).

**Fig. 8.**
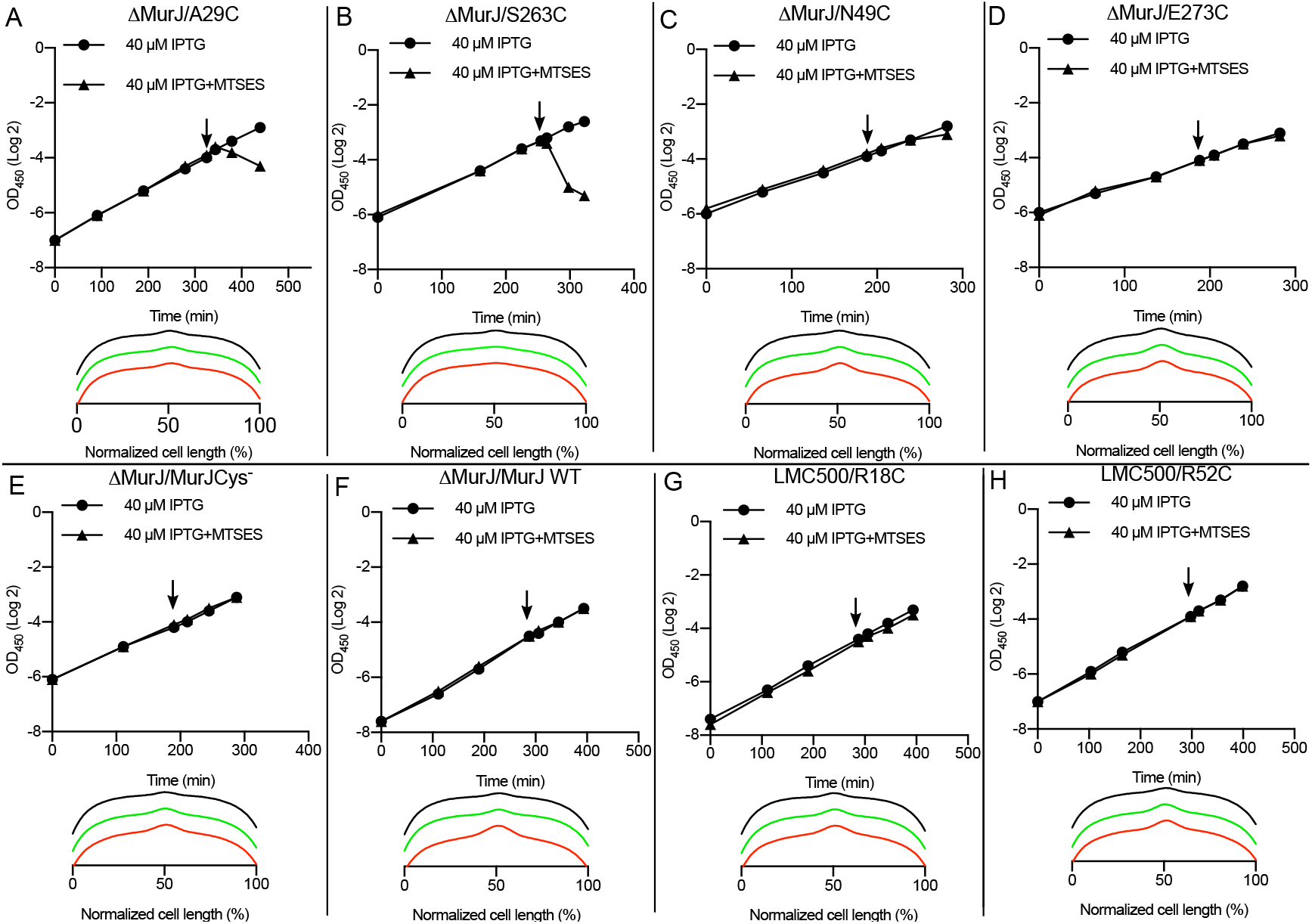
Localization of MurJ cysteine mutants in the presence of MTSES. Strains expressing MurJ single-cysteine-mutants from plasmid were grown to steady state in Gb1 medium at 28 °C, and expression was induced with IPTG for 2 MDs. MurJ localization was determined at 10 min and 40 min after the addition of MTSES (for mutant S263C, localization was determined at 10 min and 20 min because of its higher sensitivity to MTSES), Average of MurJ localization profiles were plotted along normalized cell axis. Black lines indicate the MurJ localization in the absence of MTSES, green and red lines indicate the MurJ localization at 10 and 40 min in the presence of MTSES, respectively. More than 1200 cells were included for each experiment. (A-D) Only growth but not localization of functional mutants A29C, S263C, N49C and E273C is affected by MTSES. (E) Neither growth nor localization of cysteine free mutants is influenced by MTSES. (F) Neither growth nor localization of wild type MurJ is influenced by MTSES. (G and H) Localization of non-functional MurJ mutants R18C and R52C is not affected by MTSES.

Next, we also investigated whether MTSES influences the localization of the total-loss-of-function mutants, R18C, R24C, R52C, and R270C. Similar MTSES experiments were performed on these mNG-fused mutants in the LMC500 background. To eliminate the potential influence of MTSES on wild type MurJ from the LMC500 genome, the MurJ deletion strain expressing mNG fused wild-type MurJ was firstly checked. In agreement with the evidence that the two native cysteine residues in wild-type MurJ are not labeled by MTSES (Butler et al., 2013), no defect on cell growth, morphology or MurJ localization was observed when MTSES was added (Fig. 8F and Fig. S11F). Similarly, no growth or morphology defect was observed after addition of MTSES to the non-functional single cysteine mutants, and MurJ still localized at midcell (Fig. 8G-H, Fig. S11, G-K and Fig. S12). To point out, when induced with 40 μM IPTG, mutant R24C showed strongly reduced fluorescence and a cytoplasmic-like localization, compared to other mutants. However, when induced with 100 μM IPTG, R24C showed a normal MurJ localization and comparable fluorescence, indicating that mutation R24C does not affect MurJ localization (Fig. S12C). Surprisingly, a spontaneous suppressor mutation R447H that is situated at the cytoplasmic end of TMH13 restored fluorescence and showed a typical MurJ localization when induced with 40 μM IPTG (Fig. S7D-F). Since the R24C mutant was reported not to affect protein level (Butler et al., 2014), R24C might slightly change the structure of MurJ, and affect mNG folding and fluorescence, while mutation R447H could somehow suppress this defect.

Together, the investigation of MTSES on MurJ cysteine variants supports the notion that MurJ activity is not essential for its localization.

## Discussion

### Recruitment of MurJ to midcell and the coordination of cell division and septal PG synthesis

The synthesis of PG has been studied for decades, and FtsW and RodA have been always considered as the lipid II flippases in elongasome and divisome, respectively (Dai & Lutkenhaus, 1991; Den Blaauwen et al., 2008; Ikeda et al., 1989; Matsuzawa, Hayakawa, Sato, & Imahori, 1973; Mohammadi et al., 2011, 2014; Sieger, Schubert, Donovan, & Bramkamp, 2013). Only recently, MurJ was identified as the lipid II flippase (Elhenawy et al., 2016; Meeske et al., 2015; Qiao et al., 2017; Ruiz, 2008; Sham et al., 2014). FtsW has been shown to bind and flip lipid II and other lipids *in vitro* but not *in vivo* (Mohammadi et al., 2011, 2014), whereas evidence has been provided that MurJ has the lipid II flipping activity *in vivo*, but only lipid II binding activity so far *in vitro* (Bolla et al., 2018; Leclercq et al., 2017; Qiao et al., 2017; Sham et al., 2014). Our study indicates that these observations can be very well all valid, as both FtsW and MurJ are required for lipid II flipping during cell division.

Our results show for the first time the specific MurJ cellular localization in *E. coli.* Its midcell localization is critically dependent on maturation of the divisome, on PBP3 and FtsW activity, and lipid II synthesis. Interestingly, in *E. coli*, MurJ can be replaced by non-homologous flippases from other organisms (Elhenawy et al., 2016; Fay & Dworkin, 2009; Kuk et al., 2017; Meeske et al., 2015; Ruiz, 2009). Probing the interaction of MurJ with cell division proteins such as PBP3, FtsW, MurG using our cytoplasmic FRET assay (Alexeeva, Gadella, Verheul, Verhoeven, & Den Blaauwen, 2010) did not support interaction between these proteins and MurJ, which is also supported by the loss of MurJ localization in aztreonam treated cells that still have an intact divisome at mid cell (Fig. 4). Although negative FRET results are not at all a guarantee for the absence of protein interactions, together with the smoothly replacement of MurJ by other non-homologous flippases, it suggests that protein-protein interactions are likely not involved in MurJ recruitment, which implies that its localization at midcell is driven by its substrate lipid II. This is also in agreement with the delocalization of MurJ in the presence of D-cycloserine that inhibits the production of Lipid-II (Fig. 5). Inactivation of MurJ *In vivo* results in accumulation of lipid II (Qiao et al., 2017; Ruiz, 2008), indicating that lipid II synthesis is independent of the flipping process and MurJ’s presence is needed for lipid II translocation but not synthesis. The failure of MurJ to localize at an incomplete divisome or upon inactivation of PBP3 (by aztreonam) or FtsW (by mutation) signifies that lipid II is inaccessible to MurJ under these conditions (Fig 3, 4 and 6).

But why would lipid II not be accessible to MurJ under these conditions? One possible explanation could concern the regulation between PBP3 and FtsW to bind lipid II. We showed that MurJ requires FtsW activity for midcell localization. FtsW is clearly able to bind lipid II *in vitro* (Leclercq et al., 2017), and the absence of PBP3 causes FtsW to hold lipid II and prevents its polymerization by PBP1b, whereas presence of PBP3 stimulates the release of lipid II from FtsW and allows its polymerization by PBP1b (Leclercq et al., 2017). Adding that an incomplete divisome will keep the FtsW-PBP3 complex activity in check (probably) by the FtsQLB subcomplex (Liu et al., 2015; Weiss, 2015), a logical explanation would be that the inactivated FtsW, either caused by the incomplete divisome assembly, or inactivation of PBP3, or mutations (FtsW R145A and FtsW K513N), is unable to release lipid II to MurJ, and thus blocks MurJ midcell recruitment. An alternative explanation could be that FtsW, stimulated by PBP3, is required to take over the flipped lipid II from MurJ to use it or present it for PG synthesis. The inactivated PBP3 and/or FtsW will keep MurJ from resuming its flipping cycle. In this scenario MurJ would diffuse away from the division site and would provide any available transglycosylase with lipid II. Taking over lipid II from MurJ would be in agreement with the suggested transglycosylase function of FtsW (Meeske et al., 2016). The requirement of FtsW/PBP3 activities would also explain why MurJ is not flipping lipid II *in vitro.* A third possibility, which we cannot exclude presently, is that FtsW and MurJ are both flipping lipid to insert multiple glycan stands simultaneously in the septum as was suggested in the three for one or four for two models of PG insertion during cell division (Egan & Vollmer, 2015).

Why would it be necessary to present lipid II at midcell to MurJ (or take it over from MurJ) and does this imply that RodA as an FtsW homolog is also presenting (or taking over) lipid II to/from MurJ during lateral growth? The midcell position is very precisely determined by the cell (Ortiz, Natale, Cueto, & Vicente, 2015; Rowlett & Margolin, 2015). If lipid II would be synthesized randomly and flipped randomly, it could lead to loss of precision of the binary fission into identical daughter cells. The observation that inhibition of the elongasome did not affect division or MurJ midcell localization, and the inhibition of the divisome did not influence cell length growth (Fig. 4 and Fig. S6), suggest that lipid II flipping is precisely organized at the position where new PG synthesis should occur. This can be achieved by allowing it only to flip as closely as possible where the substrate is needed, and by only allowing it to bind lipid II next to the proteins that are activated to synthesize the new cell wall.

Other studies suggested that MurJ alters between inward-facing and outward-facing conformations during substrate transport (Kuk et al., 2017), and observed that cysteine mutations at the charged residues, R18, R24, R52 and R270, in the cavity center cause the loss function of MurJ, while some other cysteine mutants at position A29, N49, S263 and E273 are functional but not in the presence of cysteine-reactive agent MTSES (Butler et al., 2014; Sham et al., 2014). But how do these residues contribute to the function of MurJ is not clear. Our results show that the functionality of MurJ is not required for its midcell recruitment, despite the inactivation caused directly by mutation or by MTSES treatment (Fig. 7 and 8). As we mentioned above, MurJ midcell recruitment is likely driven by its substrate lipid II, our results on the MurJ mutants suggest that these mutants are able to recognize at least part of the substrate lipid II, but cannot flip. Also, these important residues are not conserved in other flippases that are able to replace MurJ and therefore more likely involved in the flipping mechanism than in initial substrate recognition (Fig. S13).

However, residues A29 and S263 are situated at the interface between the two lobes of MurJ, and thus are predicted to be important for MurJ conformational changes from the outward-facing to the inward-facing state, and binding of MTSES at these two positions will trap the protein at the outward-facing state (Kuk et al., 2017; Sham et al., 2014). Also, recent *in vitro* evidence suggests that binding of MTSES at A29C reduced the lipid II binding affinity of MurJ protein (Bolla et al., 2018). This would predict delocalization for the MTSES bound A29C MurJ mutant, whereas we observe the typical midcell localization *in vivo.* If this MTSES bound mutant does not recognize lipid II, its localization would require protein-protein interaction, which is unlikely in view of MurJ’s smoothly replacement by completely non-homologous flippases. It would also imply that the dependence of MurJ recruitment on divisome assembly, lipid II synthesis, FtsW activity and PBP3 activity would then be indirect, and that an important unknown factor that determines MurJ localization is missing. *In vitro* data are not always completely comparable to the *in vivo* situation, and given the conflicting lipid II binding evidences *in vitro* (Bolla et al., 2018; Leclercq et al., 2017), we assume that the MurJ A29C MTSES mutant still has a partial ability to recognize lipid II. According to the present published models (Chamakura et al., 2017; Kuk et al., 2017; Qiao et al., 2017), this would require changing to the inward open conformation, which is the only conformation observed thus far in MurJ crystals (Kuk et al., 2017).

### Model of MurJ function in septal PG synthesis

Based on our evidence and that of others (23, 38, 39, 41, 43, 44, 90, 91), we arrived at a model for the mechanism of MurJ recruitment and lipid II flipping during septal PG synthesis (Fig. 9). After maturation of the core divisome complex, MurJ is recruited to midcell through the recognition of its substrate lipid II (Fig. 9 A), and ensures the precise positioning of septal PG synthesis. Interruption of lipid II synthesis by D-cycloserine, or inactivation of PBP3 by aztreonam, or inactivation of FtsW will likely make lipid II inaccessible for MurJ, and block the recruitment of MurJ to midcell. Once recruited, MurJ flips lipid II by changing from the inward-facing state to an outward-facing conformation, and the flipped lipid II can be used for new PG synthesis (Fig. 9 B and C). Subsequently, the unloaded outward facing MurJ switches back to the inward-facing state, and can be recruited for new cycles (Fig. 9D). Inactivation of MurJ, either by direct mutation at the charged residues R18, R24, R52, R270, or by binding of MTSES to residues A29C, N49C, S263C and E273C, will disrupt the translocation of lipid II across the membrane (Fig. 9B). Although we cannot totally rule out the possibility of involvement of protein-protein interaction recruitment of MurJ during septal PG synthesis, our data show a possible mechanism of how MurJ functions during cell division *in vivo*, and visualize of MurJ localization will give possibilities for future investigations and further antibiotics developments. It should be noted that although we mainly focused on the septal MurJ, some evidence, like the D-cycloserine experiments, MurJ depletion experiment, and MTSES experiments, showed a global effect on *E. coli*, indicating that MurJ very likely works similarly in collaboration with the elongasome, likely with the FtsW homologous protein RodA.

**Fig. 9.**
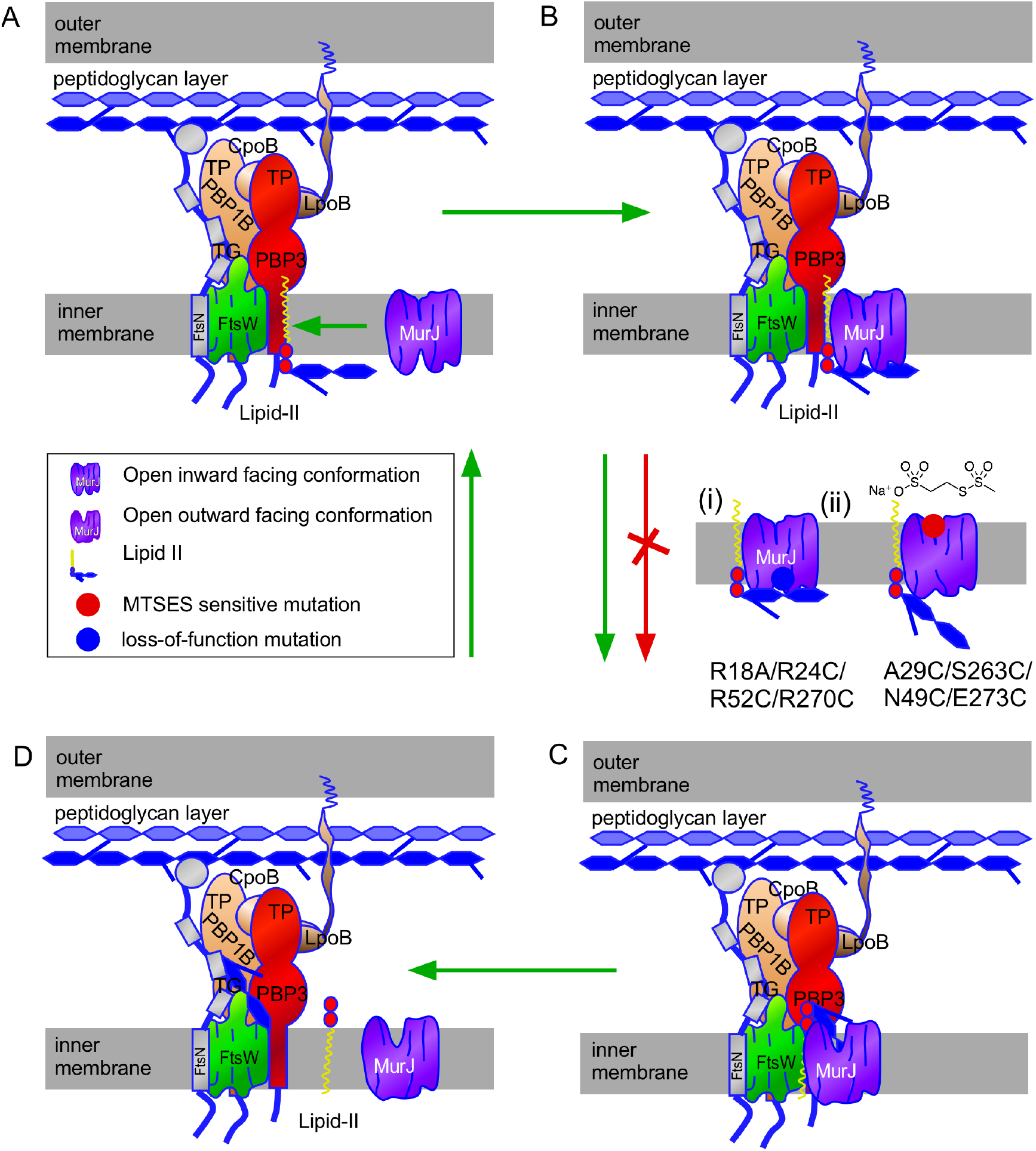
Model of MurJ recruitment and function in septal peptidoglycan synthesis. (A) Recruitment of MurJ to midcell when FtsW is activated. (B) MurJ binds substrate. (C) MurJ flips lipid II across the IM by changing its structure from the inward-facing conformation into the outward-facing state. (D) Unloaded MurJ changes back into the default inward-facing state to participate in a new cycle. (i) Loss-of-function MurJ mutants that localize but are not able to flip lipid II. (ii) Functional MurJ mutants that are inactivated by MTSES still localize but are not able to flip lipid II.

## Material and methods

### Strains and plasmids construction

Strains and plasmids used in this study are listed in Table. S1. To construct the MurJ N-terminal fluorescent protein (FP) fusion plasmids, mCherry (mCh) (Cranfill et al., 2016) or *E. coli* codon optimized mNeonGreen mNG (Meiresonne et al., 2017), were firstly cloned into plasmids pTHV037 (Den Blaauwen et al., 2003) and pSAV057 (Alexeeva et al., 2010) by restriction and ligation *(Nco*I and *Eco*RI), to generate plasmid pSAV047 and pSAV057-mNG, respectively. Subsequently, amplified MurJ fragments with different linkers were cloned into these two plasmids by restriction and ligation (*Eco*RI and *Hind*III), to generate the derived N-terminal fusion plasmids. To construct the MurJ C-terminal FP fusion plasmids, MurJ gene was firstly cloned into plasmids pTHV037 and pSAV057 with NcoI and *Bam*HI, to generate plasmids pXL09 and pXL10. Subsequently, FP genes with different linkers were cloned into these two plasmids with *Bam*HI and *Hind*III, to generate the derived C-terminal fusion plasmids. GlpT fusion plasmid pXL28 was constructed in the same way. To construct the MurJ mutant plasmids, QuickChange site directed mutagenesis (Agilent technologies, Santa Clara, CA) and Gibson assembly (Gibson et al., 2009) approaches were applied. Plasmid pXL74 was firstly generated from pXL05 by replacing the wild type MurJ with the cysteine free mutant that was amplified from plasmid pFLAGMurJΔCys (Butler et al., 2013). Single-cysteine-mutant plasmids were constructed afterwards by replacing the original residues with cysteine. The double mutation plasmid pXL93 was constructed by introducing the additional R52C mutation into plasmid pXL82.

The MurJ deletion strain XL03 (LMC500 Δ*murJ*::FRT/pRC7MurJKan) was constructed by λ-Red recombination (Datsenko & Wanner, 2000) in the presence of the complementing plasmid pRC7MurJ (Butler et al., 2013). Primers priXL17 and priXL18 were used to amplify the MurJ upstream homologous sequence region ahead of MurJ start codon from the *E. coli* genome, primers priXL19 and priXL20 were used to amplify the chloramphenicol (cat) cassette from plasmid pKD3(Datsenko & Wanner, 2000), and primers priXL21 and priXL22 were used to amplify the downstream homologous sequence that comes after MurJ stop codon from the *E. coli* genome. An overlap PCR was performed using the obtained homologous sequences and *cat* gene with primers priXL17 and priXL21 to generate the final recombination fragment. This PCR product was used to generate the MurJ depletion strain sNM01 (LMC500 ΔmurJ::cat pRC7MurJKan). Subsequently, the chloramphenicol resistance cassette was removed by plasmid pCP20 (Datsenko & Wanner, 2000), which resulted in the final *murJ* deletion strain XL03/pRC7MurJΔKan (The kanamycin cassette on this plasmid is also removable).

To prove the existence of the native *murJ* promoter, a chloramphenicol cassette was inserted after the stop codon of *yceM* upstream of *murJ* with a similar strategy as above. Primers priXL49 and priXL50 were used to amplify the upstream homologous sequence from the *E. coli* genome, priXL54 and priXL52 were used to amplify the cat cassette from plasmid pKD3, and priXL45 and priXL48 were used to amplify the downstream homologous sequence from the *E. coli* genome. The amplified overlap PCR product with primers priXL49 and priXL48 was used to construct the recombinant strain XL04 that interrupted MurJ expression from the upstream gene cluster. To construct MurJ chromosomal fusion strains that are expressed under its native promoter, plasmids pXL36 and pXL37 were firstly constructed with Gibson assembly: primers priXL51 and priXL52 were used to amplify the pKD3 backbone, primers priXL45 and priXL46 were used to amplify the *Pmurj* sequence from the *E. coli* genome, primers priXL53 and priXL47 were used to amplify FPs-MurJ sequence from plasmid pXL05 and pNM037. The final recombination fragments were generated by overlap PCR of 2 fragments with primers priXL17 and priXL48 from fragment *yceM-cat* amplified from the XL04 genome with primers priXL17 and priXL50, and fragment *cat-P_murj_-FPs-murJ* amplified from these assembled plasmids with primers priXL54 and priXL48. To construct the MurJ chromosomal FP fusion strains XL06 and XL07 expressing MurJ under control of the *P_trcdown_* promoter, fragment *yceM-cat* was amplified from the XL04 genome with primers priXL17 and priXL52, and fragment *P_trcdown_-FPs-MurJ* was amplified from plasmids pXL06 or pNM037 with primers priXL77 and priXL48. The two fragments were used for an overlap PCR with primers priXL17 and priXL48 and the resulting sequence was used to construct strains XL06 and XL07 from LMC500 by λ-Red recombination. To insert the mNG-MurJ fusion in the chromosomal of temperature sensitive strains and deletion strains, PCR fragment *yceM-cat-P_trcdown_-NG-(GGS)_2_-MurJ* was firstly amplified from the XL06 genome with primer priXL17 and priXL48, and subsequently introduced into other strains by λ-Red recombination.

To construct an FtsN depletion strain, primers priXL140 and priXL141 were used to amplify the upstream sequence of FtsN from the *E. coli* genome, priXL54 and priXL52 were used to amplify the *cat* cassette from plasmid pKD3, and priXL138 and priXL139 were used to amplify the *araC-P_BAD_* region from plasmid pJC83 (J. C. Chen & Beckwith, 2008). An overlap PCR with priXL140 and priXL138 was performed subsequently to generate the final product for recombination. The MurJ depletion strain was constructed in a similar way, primers priXL17 and priXL52 were used to amplify the *YceM-cat* region from the XL06 genome, priXL126 and priXL157 were used to amplify *araC-P_BAD_* region from plasmid pKD46, priXL156 and priXL48 were used to amplify the MurJ region from the *E. coli* genome. The final PCR product for MurJ depletion was generated by an overlap PCR with primers priXL17 and priXL48.

All PCR amplifications were performed using the DNA polymerase pfuX7 prepared in our lab as described (Nørholm, 2010), and all restriction enzymes used were purchased from New England Biolabs Inc (NEB, Ipswich, MA).

### Medium and growth conditions

LB medium (10 g of tryptone (Bacto laboratories, Australia), 10 g of NaCl (Merck, Kenilworth, NJ), 5 g of yeast extract (Duchefa, Amsterdam, The Netherlands) per liter) was used for growth in rich medium at 30 °C, 37 °C and 42 °C. The FtsW(*Ts*) strain was grown in LBΔNaCl medium (LB without NaCl and containing 0.1% glucose and 20 mg thymine per liter) as described previously (Pastoret et al., 2004). Gb1 minimal medium (6.33 g of K_2_HPO_4_ (Merck), 2.95 g of KH_2_PO_4_ (Riedel de Haen, Seelze, Germany), 1.05 g of (NH_4_)_2_SO_4_ (Sigma, St. Louis, MO), 0.10 g of MgSO_4_·7H_2_O (Roth, Karlsruhe, Germany), 0.28 mg of FeSO_4_·7H_2_O (Sigma), 7.1 mg of Ca(NO_3_)_2_·4H_2_O (Sigma), 4 mg of thiamine (Sigma), 2 mg of uracil (Sigma), 2 mg of lysine (Sigma), 2 mg of thymine (Sigma), and 0.5 % glucose (Merck) per liter, pH 7.0) was prepared for steady state growth at 28 °C as described (Van der Ploeg et al., 2013). Protein expression was induced with isopropyl β-D-1-thiogalactopyranoside (IPTG, Promega, Madison WI) or L-(+)arabinose (Sigma) at described concentrations. All antibiotics were purchased from Sigma Aldric, Working antibiotics concentrations were: 100 μg.mL^−1^ ampicillin (10 μg.mL^−1^ for *ΔtolA* and *Δpal* strains), 25 μg.mL^−1^ chloramphenicol, 50 μg.mL^−1^ kanamycin, 10 μg.mL^−1^ tetracycline (for chromosomal recombinant strains, half of the concentrations were used). Optical density (OD) was measured at 450 nm and 600 nm when grown in minimal medium and rich medium, respectively.

### Testing functionality of MurJ fusions

Functionality experiments were performed as described previously (Butler et al., 2013, 2014) using the MurJ depletion strain XL03 that expresses a functional MurJ copy from plasmid pRC7MurJΔKan. This plasmid has partitioning defects causing some of the daughter cells to lose this plasmid and die after division, unless they are complemented with a functional copy of MurJ since MurJ is essential for *E. coli*. In addition, pRC7MurJΔKan contains the *lacZ* operon and thus cells that lost the plasmid will also lose their ability to produce the blue color on 20 μg.L^−1^ X-gal (Sigma) agar plates, as our pTHV and pSAV-based plasmids do not. The functional FPs fusions to MurJ were obtained by selecting the white colonies on Ampicillin or chloramphenicol X-gal agar dish after transformation. The growth and fluorescence intensity of strains containing a fusion expressing plasmid were investigated with Synergy Mx BioTek plate reader and microscopy.

### Immunolocalization experiments

*E. coli* cells were fixed with FAGA (2.8% formaldehyde and 0.04% glutaraldehyde final concentration) for 15-20 minutes in a shaking waterbath after steady state growth in Gb1 minimal medium at 28 °C (Den Blaauwen et al., 2003). Subsequently, cells were permeabilized and immunolabeled with antibodies as described (Buddelmeijer et al., 2013). The antibodies against FtsZ, FtsN, MurG used in this study were purified as described (Aarsman et al., 2005; Mohammadi et al., 2007). Polyclonal antibodies against MraY were obtained from Rabbits that were injected with an antigenic peptide that was designed with the Epiros program (Biosiris). Based on the predicted topology by Bouhss et al (16,17) the peptide localizes in the cytoplasmic loop between TMH 1 and 2. The sequence was H2N-G54QVVRNDGPESHFS67C-COOH. The cysteine was added at the C-terminal to perform the coupling with the KLH carrier protein.

### Depletion or inactivation of of divisome proteins

Strain XL17, XL15, XL13 and XL14 that contain a MurJ FP fusion and the temperature sensitive divisome mutant proteins FtsW, FtsI, FtsQ and FtsA, respectively, were diluted 1:1000 from overnight cultures (grown at 30 °C) into fresh LB medium (LB Δ NaCl medium for XL13) with 12.5 *μ* g.mL^−1^ chloramphenicol, and grown at 30 °C to OD_600_ of approximately 0.2. Cells were diluted 1:5 into the same fresh medium that was pre-warmed at 30 °C and 42 °C, respectively, and kept growing for 2 mass doublings. To deplete FtsN, strain XL23 was diluted 1:1000 from an overnight culture (grown at 30 °C) into fresh LB medium with 12.5 *μ* g.mL^−1^ chloramphenicol and 0.5% w/v glucose, and kept growing at 30 °C to OD600 of approximately 0.2. Induced with 0.2% w/v arabinose was set as control. After growth, cells were immobilized on 1% agarose slides, and imaged live by phase contrast and epifluorescence microscopy.

### MTSES assay

Sodium (2-sulfonatoethyl) methanethiosulfonate (MTSES) was purchased from Santa Cruz Biotechnology Company. A gradient assay on MTSES concentrations (0, 0.01, 0.05, 0.1 or 0.5 mg.mL^−1^) was firstly applied on Gb1 steady state grown strain XL03 that expressed the MurJ A29C mutant. MurJ localization was determined by microscopy at 10 min and 40 min after addition of MTSES. Upon induction of expression with 20 μM IPTG for 2 MDs rapid cell lysis was observed after adding MTSES for 10 min at concentrations 0.1mM or 0.5 mM. A pattern of MurJ midcell localization was observed despite the increase of MTSES concentrations (Fig. S7A). However, we noticed that lower concentration of MTSES (0.01 mM) was not sufficient to influence the cell growth, and cells lysed too fast at higher MTSES concentrations (above 0.05 mM) to determine MurJ localization at a longer time scale. Since MTSES targets MurJ A29C by covalent binding, we suspected that the higher expression of MurJ might help to increase the resistance to MTSES. Thus, 40 μM IPTG was used for MurJ induction (2 MDs). Indeed, higher expression of A29C slightly improved the survival of cells, with a yield of better fluorescent signal, and MurJ midcell localization was able to be determined at both 10 min and 40 min after MTSES addition. In addition, 0.5 mM MTSES was sufficient to show the influence on cell growth. Thus, 0.5 mM MTSES and 40 μM IPTG inductions were used for the investigation of MurJ localization in the presence of MTSES.

### Microscope and image analysis

For localization imaging, cells were immobilized on 1.0% agarose (w/v in Gb1) pads and imaged immediately. Fluorescence microscopy was carried out either with an Olympus BX-60 fluorescence microscope equipped with a CoolSnap *fx* (Photometrics) CCD camera, a 100×/N.A. 1.35 oil objective, and software ‘ImageJ-MicroManager’, or with a Nikon Eclipse Ti microscope equipped with a C11440-22CU Hamamatsu ORCA camera, a CFI Plan Apochromat DM 100 × oil objective, an Intensilight HG 130W lamp and the NIS elements software (version 4.20.01).

Images were analyzed with Coli-Inspector supported by the ObjectJ plugin for ImageJ (version 1.49v) (Vischer et al., 2015). Briefly, the length and diameter of more than 1200 individual cells were marked and analysis in the phase contrast images. Fluorescence and phase contrast images were aligned and fluorescence background was subtracted as described (18). The fluorescence of each cell was collected in a one pixel wide bar with the length of the cell. A map of the diameter or the fluorescence localization and intensity was generated with the cells sorted according to increasing cell from left to right. Because cells were grown to steady state, the length of the cells can be directly correlated to the cell division cycle age. A collective profile is created from all cell profiles in a map. They are first resampled to a normalized cell length of 100 data points, and then averaged to a single plot, in either 1 group or more age bins. The FCplus (the extra fluorescence at mid cell in comparison to the fluorescence in the rest of the cell) and Ringfraction (FCplus/Total cellular fluorescence) profiles were generated with the “graph assistant” macro in ObjectJ (18).

**Fig. 2.**
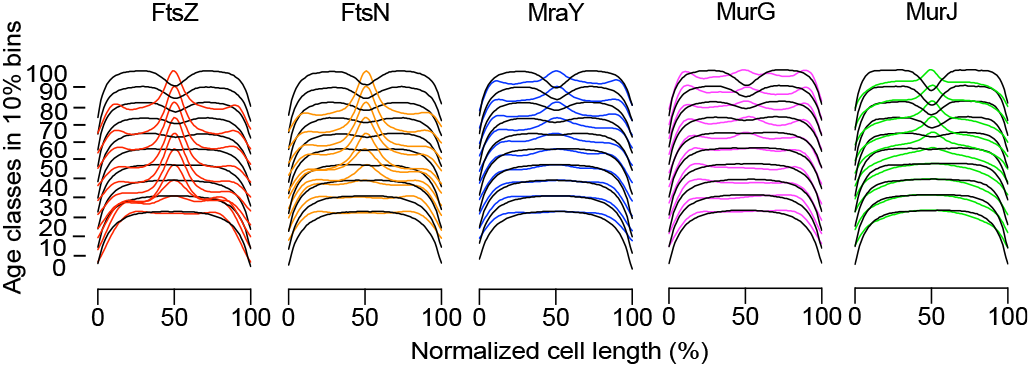
Timing of MurJ midcell localization. mNG-(GGS)_2_-MurJ cellular localization was compared with the localization of divisomal and PG precursor synthesizing proteins. XL08 strain was grown to steady state in Gb1 medium. MurJ localization was determined in living cells. After fixation with FAGA, the rest of cells were immunolabeled with antibodies against FtsZ, FtsN, MraY and MurG. Diameter (black lines) and Fluorescence (colored lines) profiles were plotted into 10 % age class bins along the normalized cell length in %. More than 1200 cells were included.

## ACKNOWLEDGEMENTS

We would like to thank Dr. N. Ruiz for critically reading the manuscript and for the gift of plasmids pFLAGMurJΔCys and pRC7MurJ for cloning and the functionality testing.

X.L. was supported by the Chinese Scholarship Council (File No.201406220123), and N.Y.M. by NWO ALW open program (822.02.019).

X.L. and T.B. designed most experiments. X.L performed most experiments and N.Y.M designed and performed some of the experiments. X.L and TdB analyzed the data and wrote the manuscript. A.B. isolated MraY and made antibodies against the protein. All authors critically read the manuscript and gave feedback.

## References

Aarsman, M. E. G., Piette, A., Fraipont, C., Vinkenvleugel, T. M. F., Nguyen-Distèche, M., & Den Blaauwen, T. (2005). Maturation of the *Escherichia coli* divisome occurs in two steps. Molecular Microbiology, 55(6), 1631–1645. https://doi.org/10.1111/j.1365-2958.2005.04502.x

Al-Dabbagh B, Henry X, El Ghachi M, Auger G, Blanot D, Parquet C, Mengin-Lecreulx D, B. A. (2008). Active Site Mapping of MraY, a Member of the Polyprenyl-phosphate N-Acetylhexosamine 1-Phosphate Transferase Superfamily, Catalyzing the First Membrane Step of Peptidoglycan Biosynthesis †. Biochemistry, 47(34), 8919–8928. https://doi.org/10.1021/bi8006274

Alexeeva, S., Gadella, T. W. J., Verheul, J., Verhoeven, G. S., & Den Blaauwen, T. (2010). Direct interactions of early and late assembling division proteins in *Escherichia coli* cells resolved by FRET. Molecular Microbiology, 77(2), 384–398. https://doi.org/10.1111/j.1365-2958.2010.07211.x

Azam, M. A., & Jayaram, U. (2015). Inhibitors of alanine racemase enzyme: a review. Journal of Enzyme Inhibition and Medicinal Chemistry, 6366(August), 1–10. https://doi.org/10.3109/14756366.2015.1050010

Barreteau, H., Kovač, A., Boniface, A., Sova, M., Gobec, S., & Blanot, D. (2008). Cytoplasmic steps of peptidoglycan biosynthesis. FEMS Microbiology Reviews, 32(2), 168–207. https://doi.org/10.1111/j.1574-6976.2008.00104.x

Bertsche, U., Kast, T., Wolf, B., Fraipont, C., Aarsman, M. E. G., Kannenberg, K., … Vollmer, W. (2006). Interaction between two murein (peptidoglycan) synthases, PBP3 and PBP1B, in *Escherichia coli.* Molecular Microbiology, 61(3), 675–690. https://doi.org/10.1111/j.1365-2958.2006.05280.x

Bisson Filho, A. W., Hsu, Y.-P., Squyres, G., Kuru, E., Wu, F., Jukes, C., … Garner, E. (2017). Treadmilling by FtsZ filaments drives peptidoglycan synthesis and bacterial cell division. Science, 355, 739–743. https://doi.org/DOI: 10.1126/science.aak9973

Blaauwen, T. Den, Buddelmeijer, N., Aarsman, M. E. G., Hameete, C. M., & Nanninga, N. (1999). Timing of FtsZ assembly in *Escherichia coli*. Journal of Bacteriology, 181(17), 5167–5175.

Bolla, J. R., Sauer, J. B., Wu, D., Mehmood, S., Allison, T. M., & Robinson, C. V. (2018). Direct observation of the influence of cardiolipin and antibiotics on lipid II binding to MurJ. Nature Chemistry, (January). https://doi.org/10.1038/nchem.2919

Bouhss, A., Mengin-Lecreulx, D., Le Beller, D., & Van Heijenoort, J. (1999). Topological analysis of the MraY protein catalysing the first membrane step of peptidoglycan synthesis. Molecular Microbiology, 34(3), 576–585. https://doi.org/10.1046/j.1365-2958.1999.01623.x

Bouhss, A., Trunkfield, A. E., Bugg, T. D. H., & Mengin-Lecreulx, D. (2008). The biosynthesis of peptidoglycan lipid-linked intermediates. FEMS Microbiology Reviews. https://doi.org/10.1111/j.1574-6976.2007.00089.x

Buddelmeijer, N., Aarsman, M., & Blaauwen, T. den. (2013). Immunolabeling of Proteins in situ in *Escherichia coli* K12 Strains. Bio-Protocol, 3(15), 1–5. https://doi.org/10.21769/BioProtoc.852

Butler, E. K., Davis, R. M., Bari, V., Nicholson, P. A., & Ruiz, N. (2013). Structure-function analysis of MurJ reveals a solvent-exposed cavity containing residues essential for peptidoglycan biogenesis in *Escherichia coli.* Journal of Bacteriology, 195(20), 4639–4649. https://doi.org/10.1128/JB.00731-13

Butler, E. K., Tan, W. B., Joseph, H., & Ruiz, N. (2014). Charge requirements of lipid II flippase activity in *Escherichia coli.* Journal of Bacteriology, 196(23), 4111–4119. https://doi.org/10.1128/JB.02172-14

Chamakura, K. R., Sham, L.-T., Davis, R. M., Min, L., Cho, H., Ruiz, N., … Young, R. (2017). A viral protein antibiotic inhibits lipid II flippase activity. Nature Microbiology. https://doi.org/10.1038/s41564-017-0023-4

Chen, J. C., & Beckwith, J. (2008). FtsQ, FtsL and FtsI require FtsK, but not FtsN, for co-localization with FtsZ during *Escherichia coli* cell division. Molecular Microbiology. https://doi.org/https://doi.org/10.1046/j.1365-2958.2001.02640.x

Chen, X., Zaro, J. L., & Shen, W. C. (2013). Fusion protein linkers: Property, design and functionality. Advanced Drug Delivery Reviews. Elsevier B.V. https://doi.org/10.1016/j.addr.2012.09.039

Contreras-Martel, C., Martins, A., Ecobichon, C., Trindade, D. M., Matteï, P.-J., Hicham, S., … Dessen, A. (2017). Molecular architecture of the PBP2–MreC core bacterial cell wall synthesis complex. Nature Communications, 8(1), 776. https://doi.org/10.1038/s41467-017-00783-2

Cranfill, P. J., Sell, B. R., Baird, M. A., Allen, J. R., Lavagnino, Z., de Gruiter, H. M., … Piston, D. W. (2016). Quantitative assessment of fluorescent proteins. Nature Methods, 13(7), 557–562. https://doi.org/10.1038/nmeth.3891

Dai, K., & Lutkenhaus, J. (1991). *ftsZ* is an essential cell division gene in *Escherichia coli.* J Bacteriol, 173(11), 3500–3506. https://doi.org/10.1046/j.1365-2958.1997.4091773.x

Datsenko, K., & Wanner, B. (2000). One-step inactivation of chromosomal genes in *Escherichia coli* K-12 using PCR products. Proc. Nat. Acad. Sci. (USA), 971242, 6640–6645. https://doi.org/https://doi.org/10.1073/pnas.120163297

de Roubin, M. R., Mengin-Lecreulx, D., & van Heijenoort, J. (1992). Peptidoglycan biosynthesis in *Escherichia coli:* variations in the metabolism of alanine and D-alanyl-D-alanine. Journal of General Microbiology, 138 Pt 8(1992), 1751–7. https://doi.org/10.1099/00221287-138-8-1751

Den Blaauwen, T., Aarsman, M. E. G., Vischer, N. O. E., & Nanninga, N. (2003). Penicillin-binding protein PBP2 of *Escherichia coli* localizes preferentially in the lateral wall and at mid-cell in comparison with the old cell pole. Molecular Microbiology, 47(2), 539–547. https://doi.org/10.1046/j.1365-2958.2003.03316.x

Den Blaauwen, T., De Pedro, M. A., Nguyen-Distèche, M., & Ayala, J. A. (2008). Morphogenesis of rod-shaped sacculi. FEMS Microbiology Reviews, 32(2), 321–344. https://doi.org/10.1111/j.1574-6976.2007.00090.x

den Blaauwen, T., Hamoen, L. W., & Levin, P. A. (2017). The divisome at 25: the road ahead. Current Opinion in Microbiology, 36, 85–94. https://doi.org/10.1016/j.mib.2017.01.007

Derouichei, R., Bénédetti, H., Lazzaroni, J. C., Lazdunski, C., & Lloubès, R. (1995). Protein complex within *Escherichia coli* inner membrane: TolA N-terminal domain interacts with TolQ and TolR proteins. Journal of Biological Chemistry, 270(19), 11078–11084. https://doi.org/10.1074/jbc.270.19.11078

Du, S., Pichoff, S., & Lutkenhaus, J. (2016). FtsEX acts on FtsA to regulate divisome assembly and activity. Proceedings of the National Academy of Sciences, 201606656. https://doi.org/10.1073/pnas.1606656113

Dubuisson, J. F., Vianney, A., & Lazzaroni, J. C. (2002). Mutational Analysis of the TolA C-Terminal Domain of *Escherichia coli* and Genetic Evidence for an Interaction between TolA and TolB. Journal of Bacteriology, 184(16), 4620–4625. https://doi.org/10.1128/JB.184.16.4620-4625.2002

Egan, A. J. F., & Vollmer, W. (2015). The stoichiometric divisome: A hypothesis. Frontiers in Microbiology, 6(MAY), 1–6. https://doi.org/10.3389/fmicb.2015.00455

Elhenawy, W., Davis, R. M., Fero, J., Salama, N. R., Felman, M. F., & Ruiz, N. (2016). The o-antigen flippase wzk can substitute for MurJ in peptidoglycan synthesis in *helicobacter pylori* and *Escherichia coli*. PLoS ONE, 11(8), 1–16. https://doi.org/10.1371/journal.pone.0161587

Fay, A., & Dworkin, J. (2009). *Bacillus subtilis* homologs of MviN (MurJ), the putative *Escherichia coli* lipid II flippase, are not essential for growth. Journal of Bacteriology, 191(19), 6020–6028. https://doi.org/10.1128/JB.00605-09

Fenton, A. K., Mortaji, L. El, Lau, D. T. C., Rudner, D. Z., & Bernhardt, T. G. (2016). CozE is a member of the MreCD complex that directs cell elongation in *Streptococcus pneumoniae*. Nature Microbiology, 2(December 2016), 16237. https://doi.org/10.1038/nmicrobiol.2016.237

Gerding, M. A., Liu, B., Bendezú, F. O., Hale, C. A., Bernhardt, T. G., & De Boer, P. A. J. (2009). Self-enhanced accumulation of FtsN at division sites and roles for other proteins with a SPOR domain (DamX, DedD, and RlpA) in *Escherichia coli* cell constriction. Journal of Bacteriology, 191(24), 7383–7401. https://doi.org/10.1128/JB.00811-09

Gibson, D. G., Young, L., Chuang, R.-Y., Venter, J. C., Hutchison, C. A., & Smith, H. O. (2009). Enzymatic assembly of DNA molecules up to several hundred kilobases. Nature Methods, 6(5), 343–345. https://doi.org/10.1038/nmeth.1318

Goehring, N. W., Gonzalez, M. D., & Beckwith, J. (2006). Premature targeting of cell division proteins to midcell reveals hierarchies of protein interactions involved in divisome assembly. Molecular Microbiology, 61(1), 33–45. https://doi.org/10.1111/j.1365-2958.2006.05206.x

Goehring, N. W., Gueiros-filho, F., & Beckwith, J. (2005). Premature targeting of a cell division protein to midcell allows dissection of divisome assembly in *Escherichia coli.* Genes & Development, 19(1), 127–137. https://doi.org/10.1101/gad.1253805

Gray, A. N., Egan, A. J. F., van’t Veer, I. L., Verheul, J., Colavin, A., Koumoutsi, A., … Vollmer, W. (2015). Coordination of peptidoglycan synthesis and outer membrane constriction during *Escherichia coli* cell division. eLife, 4(MAY), 1–29. https://doi.org/10.7554/eLife.07118

Hong, Y., Liu, M. A., & Reeves, P. R. (2018). Progress in our understanding of Wzx flippase for translocation of bacterial membrane lipid-linked oligosaccharide. Journal of Bacteriology, 200(1), 1–14. https://doi.org/10.1128/JB.00154-17

Huang, Y. (2003). Structure and Mechanism of the Glycerol-3-Phosphate Transporter from *Escherichia coli.* Science, 301(5633), 616–620. https://doi.org/10.1126/science.1087619

Ikeda, M., Sato, T., Wachi, M., Jung, H. K., Ishino, F., Kobayashi, Y., & Matsuhashi, M. (1989). Structural similarity among *Escherichia coli* FtsW and RodA proteins and *Bacillus subtilis* SpoVE protein, which function in cell division, cell elongation, and spore formation, respectively. Journal of Bacteriology, 171(11), 6375–6378. https://doi.org/10.1128/jb.171.11.6375-6378.1989

Ishino, F., Hai Kwan Jung, Ikeda, M., Doi, M., Wachi, M., & Matsuhashi, M. (1989). New mutations *fts-36, lts-33*, and *ftsW* clustered in the mra region of the *Escherichia coli* chromosome induce thermosensitive cell growth and division. Journal of Bacteriology, 171(10), 5523–5530. https://doi.org/10.1128/jb.171.10.5523-5530.1989

Khattar, M. M., Begg, K. J., & Donachie, W. D. (1994). Identification of FtsW and characterization of a new ftsW division mutant of *Escherichia coli.* Journal of Bacteriology, 176(23), 7140–7147. https://doi.org/10.1128/jb.176.23.7140-7147.1994

Kuk, A. C. Y., Mashalidis, E. H., & Lee, S. Y. (2017). Crystal structure of the MOP flippase MurJ in an inward-facing conformation. Nature Structural and Molecular Biology, 24(2), 171–176. https://doi.org/10.1038/nsmb.3346

Leclercq, S., Derouaux, A., Olatunji, S., Fraipont, C., Egan, A. J. F., Vollmer, W., … Terrak, M. (2017). Interplay between Penicillin-binding proteins and SEDS proteins promotes bacterial cell wall synthesis. Scientific Reports. https://doi.org/10.1038/srep43306

Li, G. W., Burkhardt, D., Gross, C., & Weissman, J. S. (2014). Quantifying absolute protein synthesis rates reveals principles underlying allocation of cellular resources. Cell, 157(3), 624–635. https://doi.org/10.1016/j.cell.2014.02.033

Liu, B., Persons, L., Lee, L., & de Boer, P. A. J. (2015). Roles for both FtsA and the FtsBLQ subcomplex in FtsN-stimulated cell constriction in *Escherichia coli.* Molecular Microbiology, 95(6), 945–970. https://doi.org/10.1111/mmi.12906

Lutkenhaus, J. (2009). FtsN -Trigger for septation. Journal of Bacteriology, 191(24), 7381–7382. https://doi.org/10.1128/JB.01100-09

Macheboeuf, P., Contreras-Martel, C., Job, V., Dideberg, O., & Dessen, A. (2006). Penicillin binding proteins: Key players in bacterial cell cycle and drug resistance processes. FEMS Microbiology Reviews, 30(5), 673–691. https://doi.org/10.1111/j.1574-6976.2006.00024.x

Manat, G., Roure, S., Auger, R., Bouhss, A., Barreteau, H., Mengin-Lecreulx, D., & Touzé, T. (2014). Deciphering the Metabolism of Undecaprenyl-Phosphate: The Bacterial Cell-Wall Unit Carrier at the Membrane Frontier. Microbial Drug Resistance, 20(3), 199–214. https://doi.org/10.1089/mdr.2014.0035

Matsuzawa, H., Hayakawa, K., Sato, T., & Imahori, K. (1973). Characterization and genetic analysis of a mutant of *Escherichia coli* K-12 with rounded morphology. J. Bacteriol., 115(1), 436–442.

Meeske, A. J., Riley, E. P., Robins, W. P., Uehara, T., Mekalanos, J. J., Kahne, D., … Rudner, D. Z. (2016). SEDS proteins are a widespread family of bacterial cell wall polymerases. Nature, 1–15. https://doi.org/10.1038/nature19331

Meeske, A. J., Sham, L.-T., Kimsey, H., Koo, B.-M., Gross, C. a, Bernhardt, T. G., & Rudner, D. Z. (2015). MurJ and a novel lipid II flippase are required for cell wall biogenesis in *Bacillus subtilis.* Proceedings of the National Academy of Sciences of the United States of America, 112(20), 6437–42. https://doi.org/10.1073/pnas.1504967112

Meiresonne, N. Y., van der Ploeg, R., Hink, M. A., & den Blaauwen, T. (2017). Activity-Related Conformational Changes in d,d-Carboxypeptidases Revealed by In Vivo Periplasmic Förster Resonance Energy Transfer Assay in *Escherichia coli.* mBio. https://doi.org/10.1128/mBio.01089-17

Mengin-lecreulx, D., Flouret, B., Heijenoort, J. V. A. N., S, E. R. C. N. R., Biochimie, I. De, & Paris-sud, U. (1983). Pool Levels of UDP N-Acetylglucosamine and UDP N-Acetylglucosamine-Enolpyruvate in *Escherichia coli* and Correlation with Peptidoglycan Synthesis. Journal of Bacteriology, 154(3), 1284–1290. https://doi.org/0021-9193/83/061284-07$02.00/0

Mengin-Lecreulx, D., Texier, L., Rousseau, M., & Van Heijenoort, J. (1991). The murG gene of Escherichia coli codes for the UDP-N-acetylglucosamine:N-acetylmuramyl-(pentapeptide) pyrophosphoryl-undecaprenol N-acetylglucosamine transferase involved in the membrane steps of peptidoglycan synthesis. Journal of Bacteriology, 173(15), 4625–4636. https://doi.org/10.1128/JB.173.15.4625-4636.1991

Mohammadi, T., Karczmarek, A., Crouvoisier, M., Bouhss, A., Mengin-Lecreulx, D., & Den Blaauwen, T. (2007). The essential peptidoglycan glycosyltransferase MurG forms a complex with proteins involved in lateral envelope growth as well as with proteins involved in cell division in *Escherichia coli.* Molecular Microbiology, 65(4), 1106–1121. https://doi.org/10.1111/j.1365-2958.2007.05851.x

Mohammadi, T., Sijbrandi, R., Lutters, M., Verheul, J., Martin, N. I., Den Blaauwen, T., … Breukink, E. (2014). Specificity of the transport of lipid II by FtsW in *Escherichia coli.* Journal of Biological Chemistry, 289(21), 14707–14718. https://doi.org/10.1074/jbc.M114.557371

Mohammadi, T., van Dam, V., Sijbrandi, R., Vernet, T., Zapun, A., Bouhss, A., … Breukink, E. (2011). Identification of FtsW as a transporter of lipid-linked cell wall precursors across the membrane. The EMBO Journal, 30(8), 1425–1432. https://doi.org/10.1038/emboj.2011.61

Morgenstein, R. M., Bratton, B. P., Nguyen, J. P., Ouzounov, N., Shaevitz, J. W., & Gitai, Z. (2015). RodZ links MreB to cell wall synthesis to mediate MreB rotation and robust morphogenesis. Proc Natl Acad Sci USA, 112(40), 12510–12515. https://doi.org/10.1073/pnas.1509610112

Neuhaus, F. C. (1962). The enzymatic synthesis of D-alanyl-D-alanine. I. Purification and properties of D-alanyl-D-alanine synthetase. Journal of Biological Chemistry, 237(3).

Neuhaus, F. C., & Lynch, J. L. (1972). Studies on the inhibition of D-alanyl-D-alanine synthetase by the antibiotic D-cycloserine. BIOCHEMICAL AND BIOPHYSICAL RESEARCH COMMUNICATIONS, 8(5), 377–382. https://doi.org/https://doi.org/10.1016/0006-291X(62)90011-6

Nørholm, M. H. H. (2010). A mutant Pfu DNA polymerase designed for advanced uracil-excision DNA engineering. BMC Biotechnology, 10(1), 21. https://doi.org/10.1186/1472-6750-10-21

Ortiz, C., Natale, P., Cueto, L., & Vicente, M. (2015). The keepers of the ring: Regulators of FtsZ assembly. FEMS Microbiology Reviews, 40(1), 57–67. https://doi.org/10.1093/femsre/fuv040

Oswald, F., Varadarajan, A., Lill, H., Peterman, E. J. G., & Bollen, Y. J. M. (2016). MreB-Dependent Organization of the E. coli Cytoplasmic Membrane Controls Membrane Protein Diffusion. Biophysical Journal, 110(5), 1139–1149. https://doi.org/10.1016/j.bpj.2016.01.010

Pastoret, S., Fraipont, C., Blaauwen, T. Den, Aarsman, M. E. G., Thomas, A., Brasseur, R., & Nguyen-diste, M. (2004). Functional Analysis of the Cell Division Protein FtsW of *Escherichia coli.* Journal of Bacteriology, 186(24), 8370–8379. https://doi.org/10.1128/JB.186.24.8370

Pedro, M. A. De, Quintela, J. C., Höltje, J. V, & Schwarz, H. (1997). Murein segregation in *Escherichia coli.* Microbiology, 179(9), 2823–2834. https://doi.org/doi: 10.1128/jb.179.9.2823-2834.1997

Pichoff, S., & Lutkenhaus, J. (2002). Unique and overlapping roles for ZipA and FtsA in septal ring assembly in *Escherichia coli.* EMBO Journal, 21(4), 685–693. https://doi.org/10.1093/emboj/21.4.685

Pichoff, S., & Lutkenhaus, J. (2005). Tethering the Z ring to the membrane through a conserved membrane targeting sequence in FtsA. Molecular Microbiology, 55(6), 1722–1734. https://doi.org/10.1111/j.1365-2958.2005.04522.x

Pomorski, T., & Menon, A. K. (2006). Lipid flippases and their biological functions. Cellular and Molecular Life Sciences, 63(24), 2908–2921. https://doi.org/10.1007/s00018-006-6167-7

Priyadarshini, R., De Pedro, M. A., & Young, K. D. (2007). Role of peptidoglycan amidases in the development and morphology of the division septum in *Escherichia coli.* Journal of Bacteriology, 189(14), 5334–5347. https://doi.org/10.1128/JB.00415-07

Prosser, G. a, & de Carvalho, L. P. S. (2013). Kinetic mechanism and inhibition of *Mycobacterium tuberculosis* D-alanine:D-alanine ligase by the antibiotic D-cycloserine. The FEBS Journal, 280(4), 1150–66. https://doi.org/10.1111/febs.12108

Qiao, Y., Srisuknimit, V., Rubino, F., Schaefer, K., Ruiz, N., Walker, S., & Kahne, D. (2017). Lipid II overproduction allows direct assay of transpeptidase inhibition by [beta]-lactams. Nature Chemical Biology, 13(7), 793–798. https://doi.org/10.1038/nchembio.2388

Rico, A. I., García-Ovalle, M., Mingorance, J., & Vicente, M. (2004). Role of two essential domains of *Escherichia coli* FtsA in localization and progression of the division ring. Molecular Microbiology, 53(5), 1359–1371. https://doi.org/10.1111/j.1365-2958.2004.04245.x

Rowlett, V. W., & Margolin, W. (2015). The Min system and other nucleoid-independent regulators of Z ring positioning. Frontiers in Microbiology, 6(MAY), 1–10. https://doi.org/10.3389/fmicb.2015.00478

Ruiz, N. (2008). Bioinformatics identification of MurJ (MviN) as the peptidoglycan lipid II flippase in *Escherichia coli.* Proceedings of the National Academy of Sciences of the United States of America, 105(40), 15553–7. https://doi.org/10.1073/pnas.0808352105

Ruiz, N. (2009). *Streptococcus pyogenes* YtgP (Spy-0390) complements *Escherichia coli* strains depleted of the putative peptidoglycan flippase MurJ. Antimicrobial Agents and Chemotherapy, 53(8), 3604–3605. https://doi.org/10.1128/AAC.00578-09

Ruiz, N. (2015). Lipid Flippases for Bacterial Peptidoglycan Biosynthesis. Lipid Insights, 8(Suppl 1), 21–31. https://doi.org/10.4137/LPI.S31783

Ruiz, N. (2016). Filling holes in peptidoglycan biogenesis of *Escherichia coli.* Current Opinion in Microbiology, 34, 1–6. https://doi.org/10.1016/j.mib.2016.07.010

Sauvage, E., Kerff, F., Terrak, M., Ayala, J. A., & Charlier, P. (2008). The penicillin-binding proteins: Structure and role in peptidoglycan biosynthesis. FEMS Microbiology Reviews, 32(2), 234–258. https://doi.org/10.1111/j.1574-6976.2008.00105.x

Scheffers, D. J., & Tol, M. B. (2015). LipidII: Just Another Brick in the Wall? PLoS Pathogens, 11(12), 1–12. https://doi.org/10.1371/journal.ppat.1005213

Sham, L.-T., Butler, E. K., Lebar, M. D., Kahne, D., Bernhardt, T. G., & Ruiz, N. (2014). MurJ is the flippase of lipid-linked precursors for peptidoglycan biogenesis. Science, 345(6193), 220–222. https://doi.org/10.1126/science.1254522

Shih, Y. L., Kawagishi, I., & Rothfield, L. (2005). The MreB and Min cytoskeletal-like systems play independent roles in prokaryotic polar differentiation. Molecular Microbiology, 58(4), 917–928. https://doi.org/10.1111/j.1365-2958.2005.04841.x

Sieger, B., Schubert, K., Donovan, C., & Bramkamp, M. (2013). The lipid II flippase RodA determines morphology and growth in *Corynebacterium glutamicum.* Molecular Microbiology, 90(5), 966–982. https://doi.org/10.1111/mmi.12411

Sun, Q., & Margolin, W. (1998). FtsZ Dynamics during the Division Cycle of Live *Escherichia coli* Cells, 180(8), 2050–2056.

Suzuki, H., Nishimura, Y., & Hirota, Y. (1978). On the process of cellular division in *Escherichia coli*: a series of mutants of *E. coli* altered in the penicillin-binding proteins. Proceedings of the National Academy of Sciences of the United States of America, 75(2), 664–668. https://doi.org/10.1073/pnas.75.2.664

Taschner, P. E., Huls, P. G., Pas, E., & Woldringh, C. L. (1988). Division behavior and shape changes in isogenic *ftsZ, ftsQ, ftsA, pbpB*, and *ftsE* cell division mutants of *Escherichia coli* during temperature shift experiments. Journal of Bacteriology, 170(4), 1533–1540. https://doi.org/doi: 10.1128/jb.170.4.1533-1540.1988

Tsang, M. J., & Bernhardt, T. G. (2015). Guiding divisome assembly and controlling its activity. Current Opinion in Microbiology, 24, 60–65. https://doi.org/10.1016/j.mib.2015.01.002

van der Ploeg, R., Goudelis, S. T., & den Blaauwen, T. (2015). Validation of FRET Assay for the Screening of Growth Inhibitors of *Escherichia coli* Reveals Elongasome Assembly Dynamics. International Journal of Molecular Sciences, 16(8), 17637–54. https://doi.org/10.3390/ijms160817637

Van der Ploeg, R., Verheul, J., Vischer, N. O. E., Alexeeva, S., Hoogendoorn, E., Postma, M., … Den Blaauwen, T. (2013). Colocalization and interaction between elongasome and divisome during a preparative cell division phase in *Escherichia coli.* Molecular Microbiology, 87(5), 1074–1087. https://doi.org/10.1111/mmi.12150

van Teeffelen, S., Wang, S., Furchtgott, L., Huang, K. C., Wingreen, N. S., Shaevitz, J. W., & Gitai, Z. (2011). The bacterial actin MreB rotates, and rotation depends on cell-wall assembly. Proceedings of the National Academy of Sciences, 108(38), 15822–15827. https://doi.org/10.1073/pnas.1108999108

Vischer, N. O. E., Verheul, J., Postma, M., van den Berg van Saparoea, B., Galli, E., Natale, P., … den Blaauwen, T. (2015). Cell age dependent concentration of *Escherichia coli* divisome proteins analyzed with ImageJ and ObjectJ. Frontiers in Microbiology, 6(JUN), 1–18. https://doi.org/10.3389/fmicb.2015.00586

Vollmer, W., Blanot, D., & De Pedro, M. A. (2008). Peptidoglycan structure and architecture. FEMS Microbiology Reviews, 32(2), 149–167. https://doi.org/10.1111/j.1574-6976.2007.00094.x

Vollmer, W., Blanot, D., De Pedro, M. A., Scheffers, D., Pinho, M., Nikolaidis, I., … Dessen, A. (2005). Structural and mechanistic basis of penicillin-binding protein inhibition by lactivicins. FEMS Microbiology Reviews, 30(2), 565–569. https://doi.org/10.1128/MMBR.69.4.585

Wang, E., Walsh, C., & Walsh, C. (1978). Suicide substrates for the alanine racemase of *Escherichia coli.* Biochemistry, 17(7), 1313–1321. https://doi.org/10.1021/bi00600a028

Weiss, D. S. (2015). Last but not least: New insights into how FtsN triggers constriction during *Escherichia coli* cell division. Molecular Microbiology, 95(6), 903–909. https://doi.org/10.1111/mmi.12925

White, C. L., & Gober, J. W. (2012). MreB: Pilot or passenger of cell wall synthesis? Trends in Microbiology, 20(2), 74–79. https://doi.org/10.1016/j.tim.2011.11.004

Wood, B. W. A., & Gunsalusa, I. C. (1959). D-Alanine formation; a racemase in *Streptococcus faecalis.* Journal of Biological Chemistry, 190, 403–416.

Young, K. D. (2014). A flipping cell wall ferry. Microbiology, 345(6193), 139–140. https://doi.org/10.1126/science.1256585

Yousif SY, Broome-Smith JK, S. B. (1985). Lysis of *Escherichia coli* by beta-lactam antibiotics: deletion analysis of the role of penicillin-binding proteins 1A and 1B. J Gen Microbiol., 131(10), 2839–45.

